# Preservation of stemness in high-grade serous ovarian cancer organoids requires low Wnt environment

**DOI:** 10.1101/741397

**Authors:** Karen Hoffmann, Hilmar Berger, Hagen Kulbe, Sukanija Thillainadarasan, Hans-Joachim Mollenkopf, Tomasz Zemojtel, Eliane Taube, Silvia Darb-Esfahani, Mandy Mangler, Jalid Sehouli, Radoslav Chekerov, Elena Braicu, Thomas F. Meyer, Mirjana Kessler

## Abstract

High-grade serous ovarian cancer (HGSOC) likely originates from the fallopian tube (FT) epithelium. Here, we established 15 organoid lines from HGSOC primary tumor deposits that closely match the parental tumor mutational profile and phenotype. We found that Wnt pathway activation leads to growth arrest of these cancer organoids. Moreover, active BMP signaling is almost always required for generation of HGSOC organoids, while healthy FT organoids depend on BMP suppression by Noggin. Interestingly, FT organoids modified by stable shRNA knockdown (KD) of p53, PTEN, and Retinoblastoma (RB), also require a low Wnt environment for long-term growth, while FT organoid medium triggers growth arrest. Thus, early changes in the stem cell niche environment are needed to support outgrowth of these genetically altered cells. Indeed, comparative analysis of gene expression pattern and phenotype of normal and KD organoids confirmed that depletion of tumor suppressors triggers changes in the regulation of stemness and differentiation.

## Introduction

High-grade serous ovarian cancer (HGSOC), an occult malignancy which is diagnosed in more than 230,000 women each year (Ferlay et al., 2015), represents a major clinical challenge. Aside from the difficulties in developing new lines of targeted treatments, late detection and lack of understanding of the molecular mechanisms that drive development of the disease remain major hurdles. Studies in BRCA1/2 germline mutation carriers undergoing prophylactic cancer risk-reducing surgery (Callahan et al., 2007; Leeper et al., 2002) identified distinct early malignant changes in the distal part of the FT, leading to wide-spread acceptance of the theory that the FT is the primary tissue of origin of HGSOC (Bowtell et al., 2015; Vaughan et al., 2011). These lesions, termed **S**mall **T**ubal **I**ntraepithelial **C**arcinoma (STIC), are also routinely detected in the FT epithelium of ∼50 % of patients at an advanced stage of the disease, and were shown to have the same genomic profile as the mature cancer -indicative of a clonal relationship (Kuhn et al., 2012). Nevertheless, all tumor samples genomically and transcriptionally cluster together and closely resemble tubal epithelium, irrespective of whether STICs were detected in the FT (Ducie et al., 2017). Therefore, it remains unclear how cellular transformation occurs, and most importantly, which factors are essential for the development of invasive and metastatic properties, which are necessary for the spread of malignant cells from the FT to the ovary and beyond. Of particular interest in this context is the role of mutant p53, which is almost universally detected in metastatic HGSOC cancers (Cancer Genome Atlas Research Network, 2011), but can occasionally be found in the form of p53 signatures in healthy patients with unclear clinical significance (Lee et al., 2007). Thus, it cannot be excluded that the occurrence of mutated p53 is coupled to additional, independent transformation stimuli that provide a selective advantage during the process of transformation.

The recently reported generation of organoid cultures from ovarian cancer (Kopper et al., 2019), confirmed that this 3D *in vitro* model recapitulates the major properties of the cancer found *in vivo.* While providing a comprehensive overview of the different histological subtypes and stages of the disease, almost all primary HGSOC samples used in this study were pre-exposed to neoadjuvant chemotherapy. Thus, it remains unclear how stem cell regulation is altered in the native HGSOC tumor tissue compared to healthy epithelium. Here we defined growth conditions needed to achieve long-term expansion of HGSOC organoids from primary tumor deposits. We show that the niche requirements for cultivation of cancer organoids differ markedly from the growth conditions for maintaining epithelial homeostasis in healthy FT organoids, which depend on active Wnt and Notch as well as inhibited BMP paracrine cascades as previously established (Kessler et al., 2015).

In total, we have created 15 stable organoid lines from 13 primary deposits of advanced HGSOC patients, which match the mutational and phenotypic profile of the parental tumor. Importantly, long-term growth of engineered FT organoids with simultaneous depletion of p53, PTEN and RB could be sustained only in medium adapted for patient-derived HGSOC organoids. Comparative gene expression analysis of cancer organoids and p53/PTEN/RB triple knockdowns under different growth conditions revealed common key regulatory changes in markers of stemness and differentiation. This implies that signaling cues from the stem cell environment need to change early during tumorigenesis to facilitate the growth of mutated cells, and are preserved in the advanced disease setting. Thus, this study discovers important core principles in the development of HGSOC, which have important implications for understanding early carcinogenesis and disease progression.

## Results

### Establishment of HGSOC organoid culture

In order to define conditions for long-term *in vitro* propagation of primary cancer organoids from solid HGSOC deposits, we utilized a combinatorial screening approach, using samples obtained during primary debulking surgery. To avoid potential contribution from healthy fallopian tube or ovarian surface epithelium, only tumor samples from peritoneum and omentum deposits were used. The tissue was not pre-exposed to pharmacological agents, as all but one HGSOC patient underwent radical surgery prior to chemotherapy, in line with local clinical guidelines. Overall, 15 organoid lines were established from 13 different patients, which were classified based on TNM and FIGO staging (Supplementary Table 1). The majority had cancer deposits >2 cm that had invaded organs outside the pelvis (T3c) and spread to retroperitoneal lymph nodes (N1), but had not metastasized to more distant sites such are the liver and spleen (M0) (Fig. 1A). In order to generate a reference data set for each organoid line, the parental tumor sample was divided into 3 parts for 1) confirmation of the diagnosis by an experienced pathologists using histological analysis of standard HGSOC biomarkers (Supp. Fig. 1A), 2) isolation of DNA and RNA, and 3) isolation of cells for organoid culture (Fig. 1B).

**Figure 1.**
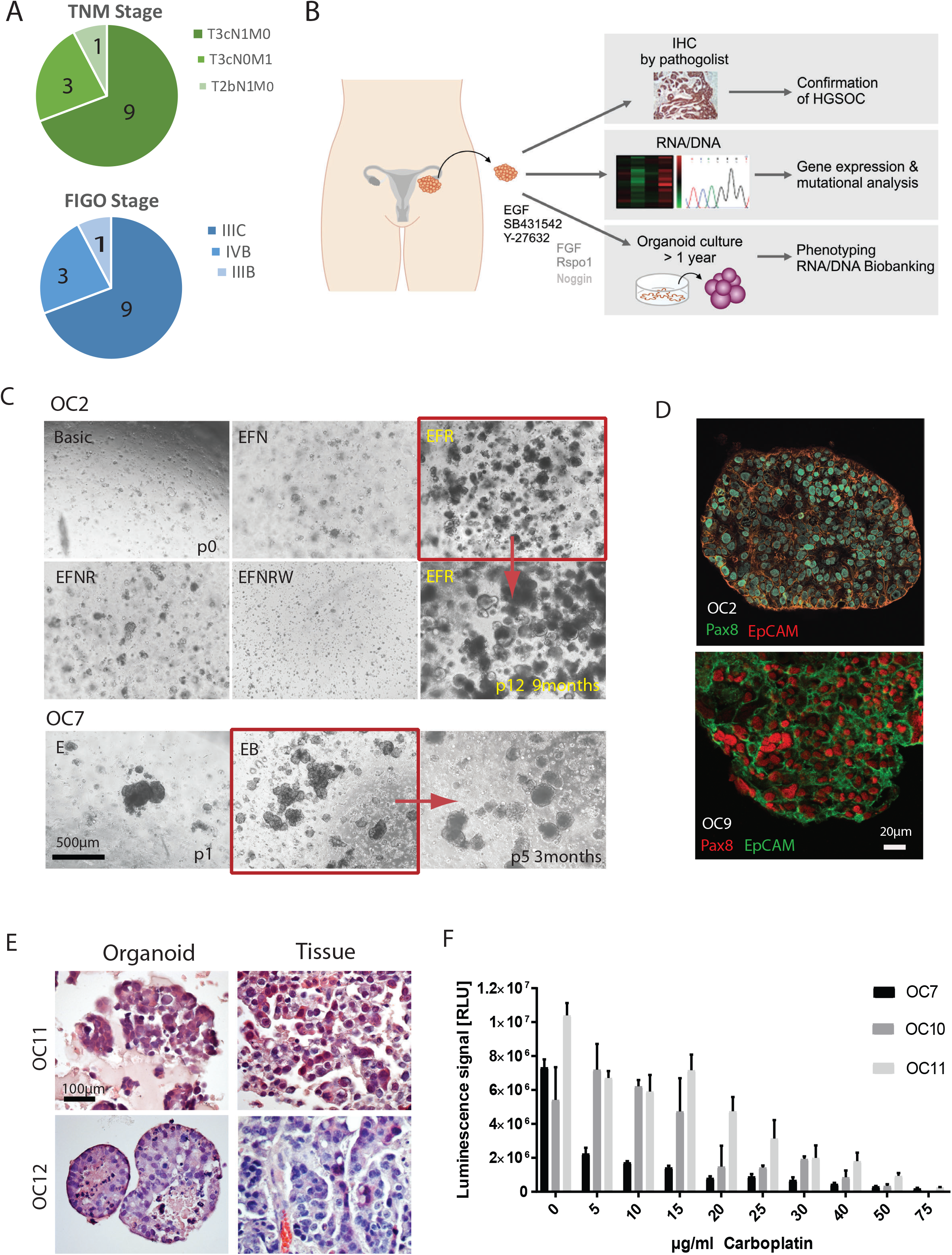
Establishment of patient-derived organoids from solid HGSOC deposits. A) Summary of cancer patient data with TNM and FIGO classifications showing advanced stage of disease at the time of surgery. B) Graphic representation of the standard experimental procedure for tumor patient material. Samples were obtained at the time of primary debulking surgery from the high purity tumor deposits in peritoneum/omentum. C) *In vitro* niche dependency of HGSOC tumor cells. Phase contrast pictures illustrate that isolated ovarian cancer cells rely on EGF supplementation for growth, while they do not grow at all in Wnt3a supplemented medium. Also, inhibition of BMP signalling through Noggin has strong negative effect on the initial growth. E – EGF, F – FGF10, N –Noggin, R – R-spondin1, B – Basic medium, P – Passage D) Cancer organoids express HGSOC markers Pax8 and EpCAM and have lost the cystic phenotype suggesting complete breakdown of epithelial polarity as seen on confocal images from two representative organoid lines. E) H&E staining of organoids and respective tissue confirm high similarity in cellular structure and tissue organization. F) HGSOC organoids show differential response to Carboplatin treatment, confirming patient-specific sensitivity of the cultures. Cell viability assay was performed after 5 days of treatment with different concentrations of Carboplatin on mature organoids from three different donors. Data represent mean ±SD of technical triplicates.

By testing media containing different combinations of growth factors (Fig. 1C) we identified conditions that support outgrowth and long-term expansion (>1 year) of cancer-derived organoids. Normal FT organoid medium (FTM) is supplemented with ROCK and TGF-β receptor inhibitors, as well as Noggin, EGF, FGF, RSPO1 and Wnt3a (Kessler et al., 2015; Kopper et al., 2019). By contrast, HGSOC organoids did not grow in this medium but required several alterations. The only paracrine growth factor that proved indispensable was EGF (E). Most notably, unlike healthy organoids from several different organs (intestinal, FT, gastric, liver etc), which require exogenous Wnt activation, not a single organoid line could be maintained in medium containing the Wnt signalling agonist Wnt3a (W) (Fig. 1C and Supplementary Table 2). Two lines did, however, expand in the presence of the Wnt agonist R-spondin 1 (RSPO1, short R), suggesting that some cultures benefit from the addition of RSPO (Fig. 1C). In addition, sustained inhibition of BMP signalling by Noggin (N) seems to be detrimental, as removal of inhibitor was a prerequisite for successful cultivation in 13/15 HGSOC samples, indicating fundamentally different roles of BMP and Wnt signalling in the regulation of stemness in HGSOC compared to normal tissue. In summary 11/15 cultures could be initiated and expanded in medium containing EGF, ROCK and TGF-β inhibitor, but without Noggin – thus allowing endogenous BMP signalling. 5 of these 11 samples benefited from the addition of BMP2 for organoid formation, further amplifying BMP signalling (B). As this patient-dependent effect of BMP2 addition was always neutral or positive, BMP2 was subsequently included in the standard medium composition. Therefore, the combination of EGF and BMP2, together with the common components of the organoid cultures ROCK, TGF-β inhibitor B27, N2 and nicotinamide was termed ovarian cancer medium (OCM).

Successfully established long-term HGSOC organoid cultures were passaged at a ratio of 1:2 to 1:3 every 10-20 days for at least 5 months prior to cryopreservation. Six lines were kept in culture for >1 year (Supplementary Table 2). Thawed cancer organoids could routinely be expanded to a multi-well screening format, and are thus suited for generating live biobanks of HGSOC organoids to explore individual therapeutic options *in vitro*.

Immunofluorescence labelling confirmed that the organoids exhibit all hallmarks of an HGSOC phenotype, including disorganized tissue architecture and loss of polarity (as marked by the absence of a central cavity), an epithelial secretory identity of all cells (strong EpCAM and PAX8 expression), and pleomorphic nuclei (Fig. 1D). HE staining further confirmed their morphological similarity to the matching tissue samples (Fig. 1E). Moreover, as proof-of-concept of the applicability of HGSOC organoids for translational research, we tested the *in vitro* drug response to carboplatin, the major first-line chemotherapeutic agent for HGSOC. As expected, organoids underwent cell death in a concentration-dependent manner, with prominent differences between organoids from different donors confirming individual variation in drug response among patients (Fig. 1F and Supp. Fig. 1B).

### HGSOC organoids match tumor tissue in mutational profile and expression of biomarkers

To test whether patient-derived organoid cultures correspond to the individual mutational profile of the parental tumor, we performed targeted sequencing for 121 candidate genes that were selected on the basis of previously published studies of ovarian cancer genomic profiles (Supplementary Table 3) (Cancer Genome Atlas Research Network, 2011; Norquist et al., 2018). Mutational analysis of 10 paired tumor fragments and organoids from 9 different patients revealed that despite the long-term expansion in vitro, they retained a high level of similarity, even including allelic frequency (Fig. 2A and Supplementary Table 4). *TP53* mutations were detected in the vast majority of samples (9/10), in agreement with the almost universal occurrence of mutated p53 in HGSOC patients (Cancer Genome Atlas Research Network, 2011). All of the *TP53* mutations in the organoid cultures were homozygous (<0.9 allele frequency), yet diverse with respect to mutation type (missense, nonsense, frame-shift or splice variant mutations). In addition to somatic mutations with proven tumorigenic potential, a number of known variant alleles were identified, including *ROCK2* (c.1292C>A), *KIT* (c.1621A>C), *MLH1* (c.655A>G) and *MSH6* (c.472C>T) which could potentially influence malignant phenotype (Brahmi et al., 2015; Kalender et al., 2010; Nakamura et al., 2014). Apart from *TP53*, *MLH1* and *MSH6*, we also found point mutations in other genes with functions in DNA repair and chromosome stability, including ATM (c.4534G>A), ATR (c.1517A>G), BRIP1 (c.254T>A) and FANCA (c.1238G>T). Interestingly, while no classic germline or somatic BRCA1/2 mutations were detected apart from three polymorphisms that are weakly associated with ovarian cancer (c.4900A>G, c.3113A>G, c.2612C>T), analysis of protein lysates revealed a significant reduction compared to healthy FT organoids in 3 of 8 cases (OC4, OC2 and OC3; Suppl. Fig. 2A). This strongly suggests the existence of epigenetic, or post-transcriptional mechanisms of BRCA1 inactivation in these cases. Altogether, 7/9 patients had mutations in one or more DNA repair gene, in congruence with the fact that HGSOC is a cancer characterized by a particularly high incidence of genomic instability. Given our finding that HGSOC organoids could not be maintained in the presence of Wnt, we noted with interest that 3 missense mutations in Wnt pathway genes (Fzd9, LRP5, TCF7L2) were identified in 3 different patients (Fig. 2A), which could impair signal transduction and thus provide further evidence that changes in this pathway play an important role in ovarian carcinogenesis. Complementary to the sequencing data, WB and IF staining were performed to examine the mutational status and subcellular localization of p53 in the organoids (Supplementary Table 4). While the nonsense mutation in OC11 as well as the splice variant and frame-shift mutations in OC4 and OC7, respectively, result in loss of p53 (Fig. 2B and Supplementary Table 5), the missense mutations in OC9 (p.R273H), OC10 (p.R175H), OC1 (p.V25F) and OC6 (p.V173M) result in a gain-of-function and nuclear accumulation phenotype with R273H and R175H being two of the most common mutations in ovarian cancer (Zhang et al., 2016) (Fig. 2C and Suppl. Fig. 2B).

**Figure 2.**
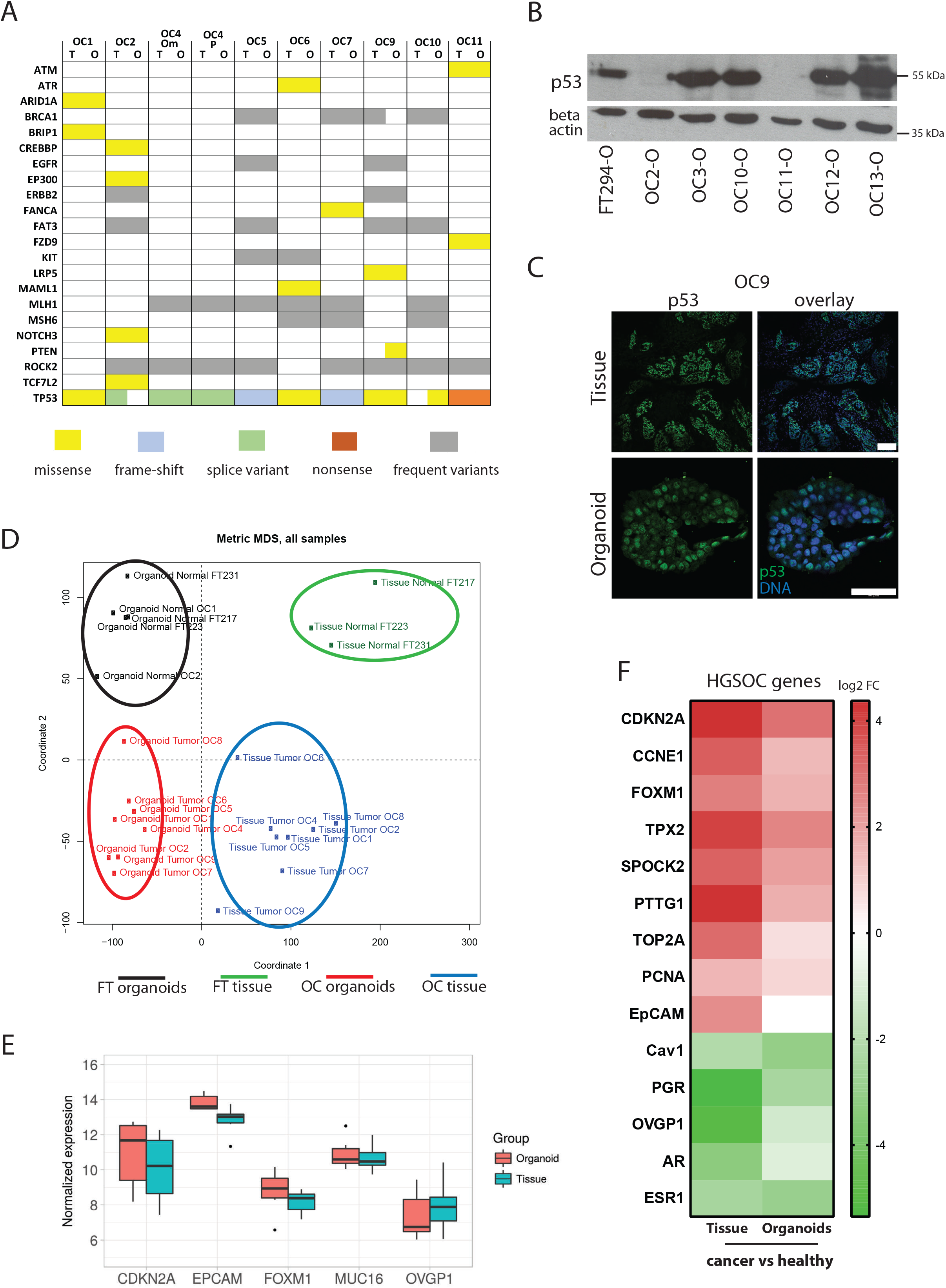
HGSOC organoids match tumor tissue in mutational profile and gene expression biomarkers. A) Overview of mutations found in organoid cultures and matching tumor tissue obtained by targeted sequencing of >200 HGSOC related genes confirms almost identical profile between parental tissue and *in vitro* long-term organoid culture. Color code indicates type of mutations which were detected. B) Representative western blot showing loss or overexpression of p53 in individual organoid lines C) Strong nuclear p53 signal in IF stainings of HGSOC organoids is indicative of a gain-of-function *TP53* mutation. Scale bars: 100 µm (Tissue) and 50 µm (Organoid). D) Multi-Dimensional-Scaling (MDS) plot based on the gene expression profiles (microarrays) of 3 healthy FT and 8 tumor tissue samples and their respective organoid cultures shows 4 clusters: Normal FT tissue, normal FT organoids, cancer organoids and cancer tissue. E) Box plots depicting overall constant level in normalized expression of major HGSOC marker genes between organoids and parental tissue. Data represent the median, quartiles, maximum, and minimum of the normalized expression from 8 different donors. F) Heat map of differentially expressed genes between cancer and healthy tissue/organoids reveals upregulation of several HGSOC biomarkers and reduction of FT differentiation markers in the cancer samples. Differential expression determined by single-color microarray for 8 different patient samples was significant for all genes with p < 0.05 except for OVGP1 and TOP2A in organoids.

To more thoroughly evaluate parallels between tissue and organoid cultures, a global gene expression analysis was performed from 8 different tumor/organoid pairs as well as 3 healthy FT tissue/organoid pairs and 2 FT organoids derived from cancer patients (also classified as normal healthy tissue). The generation of a multi-dimensional-scaling plot (MDS) revealed that the distance between OC and FT organoids is smaller than between corresponding tissues, likely due to the greater complexity of tissue samples, which contain mesenchymal and endothelial components (Fig. 2D). Still, despite the tissue heterogeneity, the respective OC organoids express very similar levels of the main cancer markers, like CDKN2A (p16), Muc16 and EpCAM, when compared to the parental sample (Fig. 2E). Moreover, correlation analysis of FT and OC organoids (Supp. Fig. 2B) shows that FT samples form a homogenous cluster and show less variability in gene expression between each other than OC organoids, which instead reflect the phenotypic diversity of ovarian tumors.

A thorough analysis of gene expression revealed that numerous hallmark genes known to be de-regulated in HGSOC are differentially expressed between OC and normal FT samples, not only in the tissue but also in organoids (Fig. 2F). This includes significant up-regulation of the cell cycle regulators CDKN2A and Cyclin E1 (CCNE1) and the transcription factor FOXM1, as well as down-regulation of differentiation markers like PGR and OVGP1.

The strong up-regulation of CCNE1 was validated by western blot analysis in 5 out of 8 samples tested (Supp. Fig. 2C and D), while RB protein itself, which is phosphorylated by CCNE1 to promote G1/S progression, was not differentially expressed (data not shown). This indicates potential RB pathway disruption by increased CCNE1 levels. Amplification of CCNE1 gene is common in HGSOC (∼20-30% of cases) but increased levels could also be a result of alternative signalling perturbations.

Together these results demonstrate that the patient-derived OC organoid lines are valid *in vitro* models of the parental HGSOC tumors, resembling not only the tissue architecture but also the mutational profile and overall gene expression.

### Stable triple knockdown of p53, PTEN and RB in FT organoids

In the next step, we wanted to analyze at which stage of disease development the observed changes in niche factor requirements occur. This is a question of particular importance in HGSOC, where transformation appears to occur in the FT but cancers are very rarely detected at this site. To mimic the cellular events, like mutation of p53 and loss of PTEN, which characterize the early stages of malignant transformation *in vitro*, we used organoids from healthy FT donor epithelium. In contrast to previous tumorigenesis models based on the transformation of immortalized primary FT monolayers (Jazaeri et al., 2011; Nakamura et al., 2018), organoids are genomically unaltered and have preserved epithelial polarity and integrity, thus ensuring mucosal homeostasis.

In order to model HGSOC development, shRNAs against p53, PTEN, and RB - the major known tumour drivers of the disease - were introduced into healthy human FT epithelial cells from donor tissues obtained during surgeries for benign gynecological conditions. The shRNAs were delivered sequentially by retro- and lentiviral vectors into epithelial cells grown in 2D culture, selected by FACS (shPTEN-mCherry, shRB–GFP) and subsequently transferred to Matrigel to initiate organoid formation (Fig. 3A and Suppl. Fig. 3A). Among the double-positive mCherry/GFP population (shPTEN, shRB), p53 depleted organoids were selected with puromycin. Successful knockdown of all 3 target genes was confirmed on RNA as well as protein level (Fig. 3B). In total 7 triple knockdown (KD) cultures were successfully generated from different FT donor tissue confirming robustness of the methodology (Suppl. Fig. 3D). In congruence with the downregulation of RB and PTEN protein, functional analysis of triple KD organoids revealed an increase in expression of CCNE1 (Fig. 3C) and elevated levels of activated phosphorylated Akt (pAkt), respectively (Suppl. Fig. 3B). Depletion of p53 protein was confirmed by resistance of organoids to the MDM2 inhibitor Nutlin3A, which triggers apoptosis in p53 WT cells (Suppl. Fig.3C).

**Figure 3.**
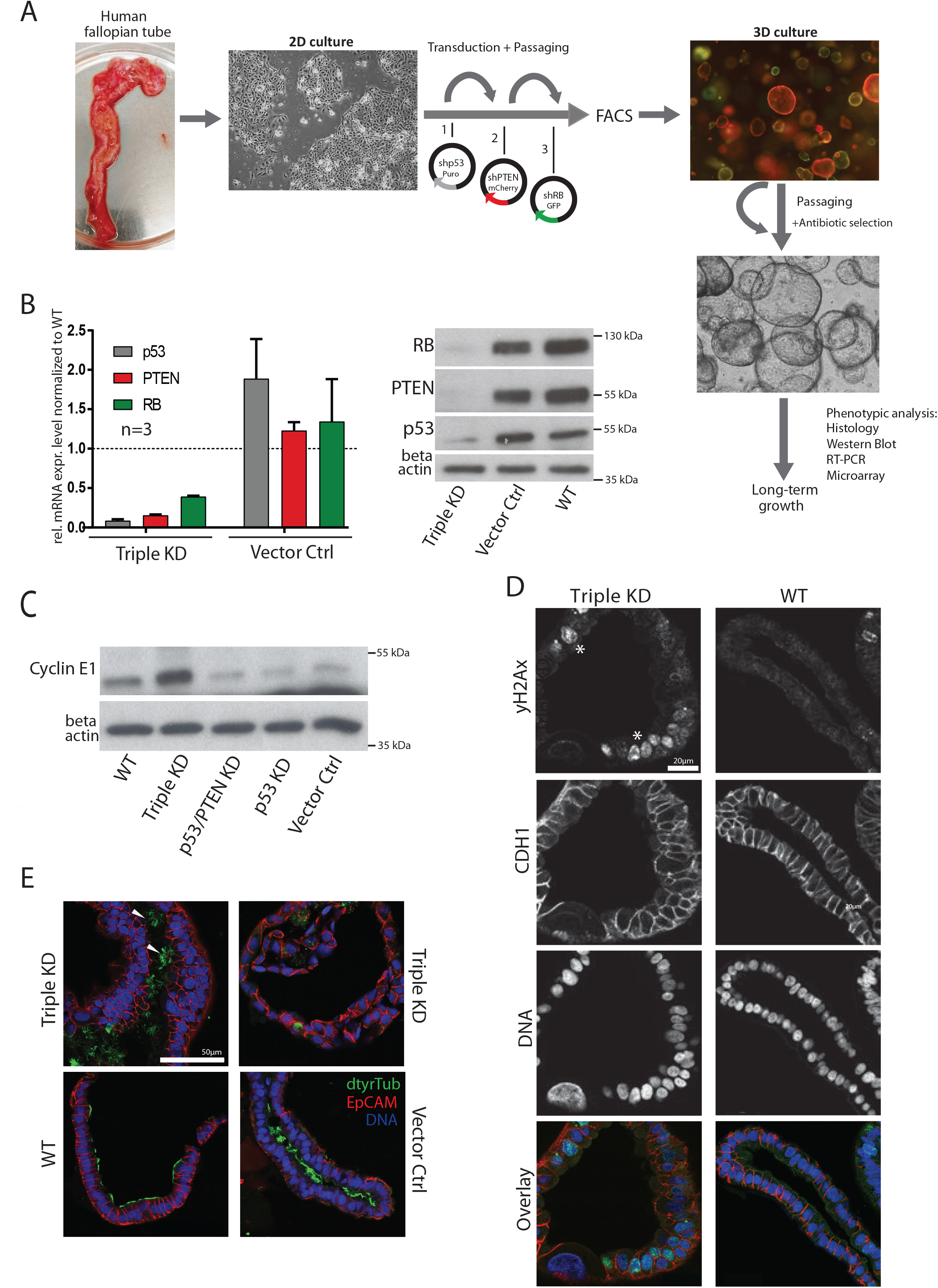
Stable triple knockdowns of tumor suppressors p53, PTEN and RB in healthy FT organoids. A) Experimental approach for genetic manipulation of FT epithelial cells (FTECs). FTECs were sequentially transduced in 2D culture with replication-deficient viruses containing specific shRNAs and different selection markers. After sorting cells were seeded in Matrigel for organoid formation. B) Confirmation of robust knockdowns of p53, PTEN and RB in FT organoids on RNA and protein level. Relative mRNA levels were normalized to the WT mRNA level and are given as the mean ±SEM from 3 different donors. A representative western blot from one donor is depicted. RNA as well as protein samples were taken at passage 1 or 2 of each organoid line. KD – Knockdown, Vec Ctrl – Vector Control, WT-Wild type C) Cyclin E1 (CCNE1) overexpression as a downstream effect of RB knockdown was confirmed by western blot analysis. D) Confocal images of triple KD organoids reveal increased DNA damage (yH2AX, marked by asterisks), atypic nuclei (Draq5) and loss of apicobasal polarity. Scale bar: 20µm. E) Presence of ciliated cells (arrowheads), as revealed by IF staining against detyrosinated (dtyr) Tubulin in triple KD organoids, proves capacity for terminal differentiation. Scale bar: 50µm.

We next compared phenotypes of KD and WT organoids by confocal microscopy. We observed loss of cell shape and misalignment of the nuclei, indicative of changes in the maintenance of apico-basal polarity – although the organization of the monolayer remained intact and the lumen was preserved with a cystic growth of the organoids. Triple KD cells also showed a stronger signal for the DNA damage marker yH2AX, indicating an increased frequency of DNA double-strand breaks and thus suggesting genomic instability (Fig. 3D). Notably, KD organoids had enlarged and polymorphic nuclei, which is one of the prominent morphological characteristics of HGSOC cells (Fig. 3D). While all these phenotypic changes suggest that the knockdown of p53, PTEN, and RB in FT organoids is pro-carcinogenic, none is sufficient as bona fide evidence of malignancy. Indeed KD cells remained competent to undergo differentiation into ciliated cells, which are thought to be the terminally differentiated cells of the FT (Fig. 3E).

### Rescue of KD organoid lines in ovarian cancer medium

Despite an initial growth advantage until the first passage after seeding, KD organoids cannot be maintained in long-term culture and undergo growth arrest after 4-8 passages, as shown by the growth curve in comparison to WT and vector control (Fig. 4A and Suppl. Fig. 4A). While individual differences among different donor cultures were observed, the premature growth arrest of KD organoids was confirmed in 7/7 cases (Suppl. Fig. 3D). This suggests that the KD organoids may undergo similar changes in stemness regulation as those observed in the HGSOC organoids.

**Figure 4.**
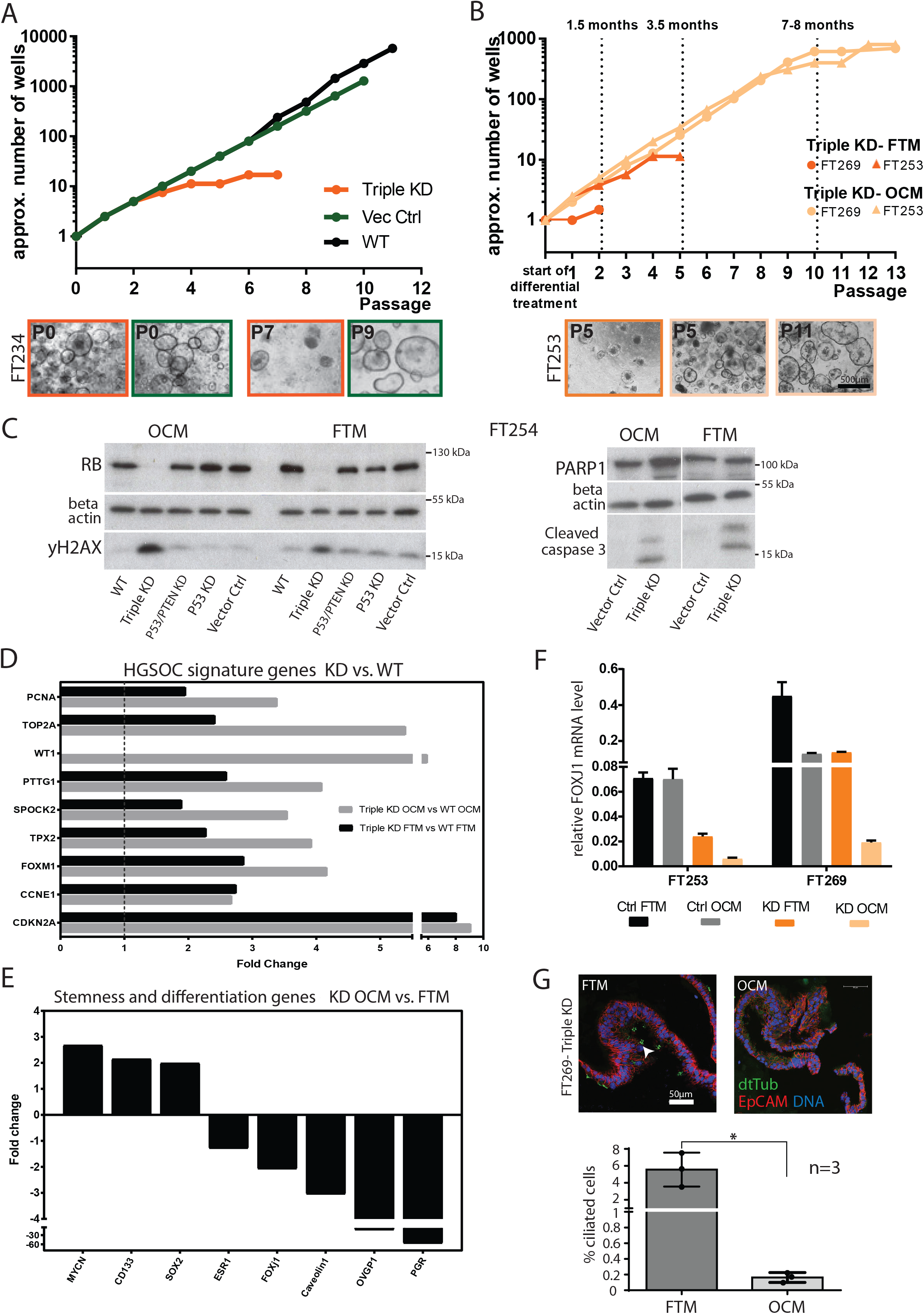
Triple KD organoids show improved growth in ovarian cancer medium. A) In long-term 3D culture triple KD organoids show loss of growth capacity and premature growth arrest compared to the controls as indicated by the representative growth curves of FT234. B) Growth curves and phase-contrast images of triple KD organoids grown in OCM compared to FTM reveal successful rescue of long-term growth capacity. C) Levels of DNA damage (yH2AX) and apoptosis (cleaved caspase-3, PARP1) do not differ among the different media as shown by representative western blots comparing triple KD organoids and their controls. D) Triple KD (p53/PTEN/RB) organoids are characterized by a HGSOC gene expression signature as revealed by microarray analysis comparing WT and KD organoids grown in either OCM or FTM. Differential expression determined by dual-color microarray for 2 biological replicates was significant for all genes with p < 0.00005. E) Differential expression of stemness- and differentiation-related genes of triple KD organoids grown in OCM vs. FTM. Fold-changes were derived from microarray data and indicate and an increase in stemness (CD133, SOX2, MYCN) and a drop in differentiation under OCM conditions (OVGP1, PGR, FOXJ1). Differential expression determined by dual-color microarray for 2 biological replicates was significant for all genes with p < 0.05. F) FOXJ1 expression determined by qRT-PCR was significantly diminished in triple KD organoids grown in OCM compared to FTM, suggesting a decrease in differentiation capacity. Data represent the mean ±SEM from technical triplicates for two independent knockdown cultures. G) Proportion of ciliated cells is significantly reduced in KD organoids under OCM growth conditions, confirming inhibition of differentiation as illustrated by confocal images. The quantification plot depicts the mean ±SEM from 3 independent biological replicates, based on quantification of the average number of ciliated cells per counted nuclei. 1000 cells were counted per individual experiment. *p < 0.05, two-sided Student’s t-test.

Indeed, growing the organoids in ovarian cancer medium improved organoid formation efficiency and enabled long-term growth (Fig. 4B). Importantly, long-term growth could only be rescued if OCM was added at passage 0/1, suggesting that inadequate niche conditions lead to irreversible loss of stemness. Phase contrast images (Suppl. Fig. 4B) confirm that while individual organoids grow to a similar size, organoid numbers are lower at each passage in FTM, finally resulting in premature growth arrest. Western blot analysis of γH2AX and cleaved caspase 3, as well as PARP1, indicated that the growth-suppressive effect of Wnt3a did not result from direct cytotoxicity, as DNA damage and apoptosis was not reduced when organoids were grown in OCM (Fig. 4C and Supp. Fig. 4C). Single p53 or double p53/PTEN KD on the other hand did not exhibit increased γH2AX levels, irrespective of the medium used, indicating that the enhanced DNA damage observed in triple KDs is the result of RB depletion.

In order to discover factors induced by culture in OCM that contribute to maintenance of stemness and longevity of the triple KD and HGSOC organoids, we performed global gene expression analysis. As expected, OCM medium devoid of Wnt and RSPO1 led to a reduction in the expression of Wnt target genes in both organoids (Suppl. Fig. 4D). The microarray data also reveal that despite cell growth arrest of the KD organoids in FTM, they continue to show elevated expression of classical HGSOC markers, including FOXM1, TOP2A, CDKN2A, and WT1, albeit at lower levels than when grown in OCM (Fig. 4D). This finding proves that knockdown of key tumor suppressor genes is sufficient to induce a degree of cellular transformation towards a cancer phenotype and supports the view that the FT epithelium is the tissue of origin of HGSOC.

However, a clear increase in the expression of genes related to stemness (CD133, MycN and SOX2) and a decrease in genes related to differentiation (PGR, OVGP1, and FOXJ1) was found in KD organoids grown in OCM vs. FTM (Fig. 4E). While PGR and ESR1 are differentiation markers connected to hormone signaling, FOXJ1 is the major transcription factor required for ciliogenesis and thus development of motile ciliated cells, presumed to be the terminally differentiated cell type in the FT mucosa. The downregulation of FOXJ1 compared to the vector control was validated by RT-PCR for organoids from 2 patients (Fig. 4F). In addition, quantification of immunolabelling against dtTubulin (Fig. 4G) in 3 independent organoid lines confirmed a lack of ciliated cells in the triple KD organoids grown in OCM, but not in FTM. Since FT mucosa renewal is driven by the differentiation of bipotent progenitors (Ghosh et al., 2017; Kessler et al., 2015), the absence of ciliated cells is suggestive of a shift towards the secretory phenotype. Notably, an outgrowth of secretory cells is thought to be a critical step in the development of HGSOC from FT epithelium. Our findings indicate that this process can be triggered *in vitro* by p53/PTEN/RB depletion, provided that canonical Wnt pathway activation is reduced.

### Wnt free medium supports stemness of HGSOC and KD organoids

Next, we wanted to confirm independently that Wnt-free medium increases stemness of KD organoids. FACS analysis of the bona fide OC stem cell marker CD133 (Kryczek et al., 2012), which is expressed on the cell surface, revealed that the number of positive cells in triple KD organoids increased when cultured in OCM vs FTM (Fig. 5A and Supp. Fig. 5A). In vector ctrl and WT cells, by contrast, the number of CD133+ cells was lower in OCM, consistent with FTM providing more optimal conditions for the preservation of stemness in healthy epithelial cells. Loss of stemness coincides with a reduction in growth potential as quantified by a luciferase-based assay determining the number of live cells after two passages of parallel cultivation in the different media (Fig. 5B). This confirms that the presence of Wnt agonists in the environment hampers the growth of p53/PTEN/RB KD organoids by repressing stemness – indicative of a substantial change in stemness regulation.

**Figure 5.**
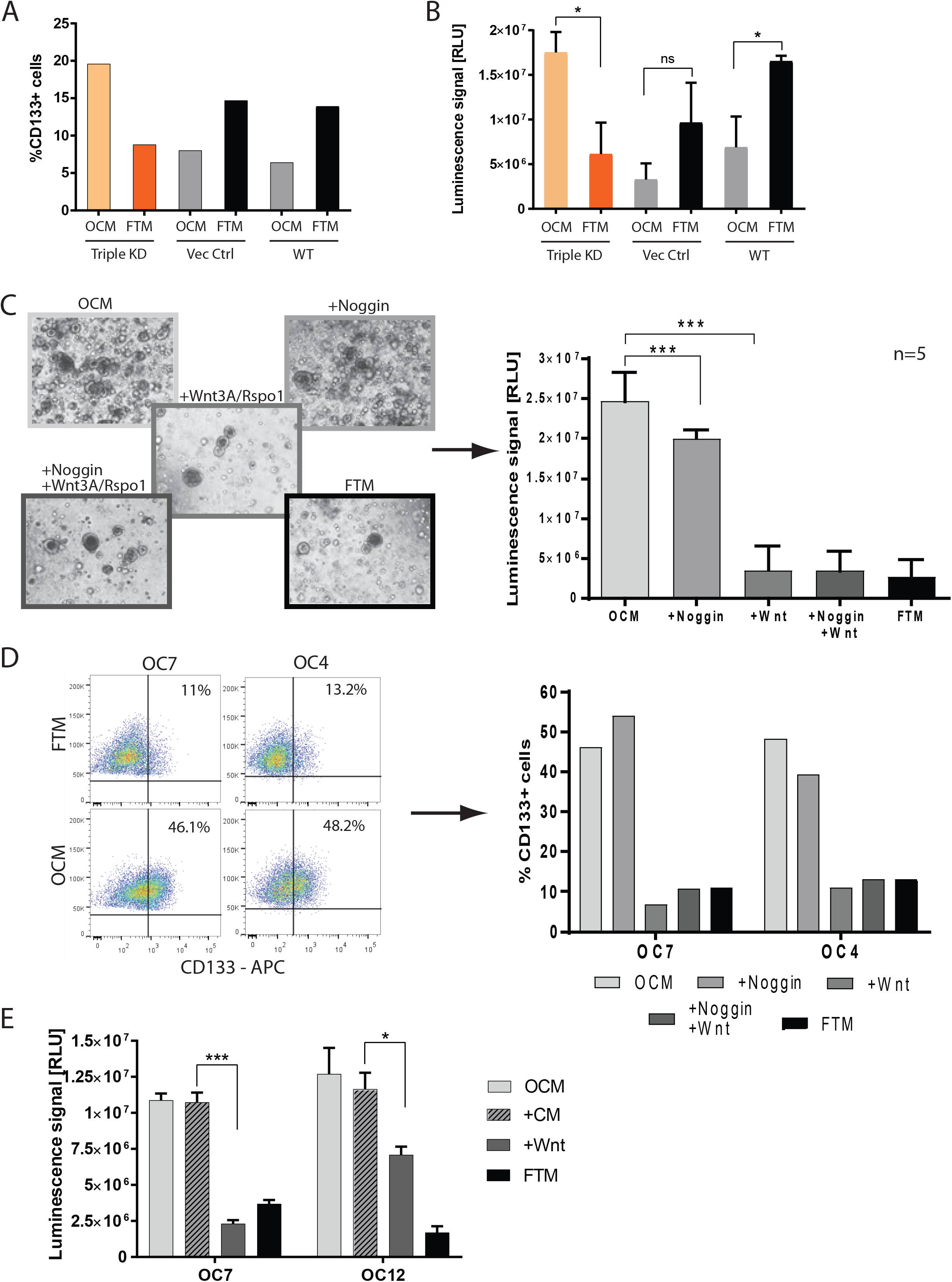
Wnt depletion supports preservation of stemness in HGSOC organoids. A) The proportion of CD133+ cells was determined by FACS analysis after growing triple KD and control cells in FTM vs. OCM for two passages. The graph is representative of two biological replicates. B) Growth capacity as well as expression of stemness markers increase when KD organoids are grown in OCM medium as shown by difference in total viable cell number (representative graph of 3 independent experiments). The luminescent cell viability assay was performed in triplicate and the bar plot depicts the mean ±SEM. * p < 0.05, two-sided Student’s t-test; KD – Knockdown, Vec Ctrl – Vector Control, WT – Wild type C) Phase contrast images of cancer organoids which entered growth arrest upon treatment with Wnt agonists over 2 passages (Scale bar: 500µm). Differences in cell number were confirmed by the respective cell viability assays (performed in technical triplicates). Data represent the mean ±SEM for 5 different OC organoid lines (n=5). **** p < 0.0001, two-sided Student’s t-test. D) The number of CD133+ cells determined by FACS for OC7 and OC4 organoids grown in different media with or without Wnt3A/Rspo1 and with or without Noggin over 2 passages, confirms a sharp drop in stemness in the presence of Wnt agonists. E) Addition of conditioned medium collected from the parental Wnt3a cell line did not suppress growth of cancer organoids, proving that the growth inhibitory effect is due to the presence of Wnt agonists. The bar plot represents the mean ±SEM of technical triplicates performed with a luminescent cell viability assay. * p < 0.05, **** p < 0.0001, two-sided Student’s t-test.

To substantiate these results, we analyzed the direct effect of Wnt3a/RSPO1 supplementation on patient-derived HGSOC organoid growth and CD133 expression. Differential treatment of cancer organoids over 2 passages clearly showed that addition of these Wnt agonists to OCM resulted in a highly significant (p < 0.0001) inhibition of organoid growth, an effect confirmed for all 6 OC organoid lines tested (Fig. 5C). Interestingly, the inhibitory effect of noggin on already established cultures was mild, but significant, confirming that BMP pathway activity is beneficial for growth but not as essential as for initial organoid formation. However, addition of noggin to mature organoid lines had no effect on the number of CD133+ cells, while Wnt3a/RSPO1 reduced numbers significantly (Fig. 5D and Supp. Fig. 5B), strongly suggesting that canonical Wnt pathway activation negatively regulates stemness in HGSOC organoids via CD133 expression.

As Wnt3a supplementation relies on the addition of conditioned medium from a Wnt-producing cell line, we next wanted to confirm that its effect was specific to the presence of Wnt. Indeed, Wnt-conditioned medium strongly upregulated the established Wnt target gene AXIN2 (Suppl. Fig. 5C). In addition, medium conditioned by the parental cell line without the Wnt-producing vector (CM) did not negatively affect organoid growth (Fig. 5E).

Overall, it can be concluded that the presence of Wnt ligands, which induce a signature of canonical Wnt target gene expression in KD and HGSOC organoids, results in a striking decrease of stemness and growth capacity in these cultures.

### MYCN and Wnt inhibitors are upregulated in HGSOC and KD FT organoids

The data from HGSOC organoids provide strong evidence that the mechanisms by which growth of cancer organoid stem cells is maintained are altered from those in the healthy epithelium and our experiments with KD organoids further show that depletion of only three tumor-driver genes can recapitulate this effect. To get a better insight into putative candidate genes that could contribute to the maintenance of stemness in cancer organoids, we compared microarray data of healthy FT epithelium and HGSOC tissue as well as patient-derived cancer and KD organoids grown in OC vs FT medium (Fig. 6A). The proto-oncogene MYCN was uniformly upregulated in cancer tissue, as well as triple KD organoids cultured in OCM, suggesting that it may play a role in driving stemness and cancer growth in HGSOC. qPCR confirmed MYCN expression to be significantly higher in cancer organoids compared to FT organoids (Fig. 6B, left panel). In addition, expression levels of MYCN relative to GAPDH were validated by qPCR in 3 independent biological replicates of triple KD organoids cultured in OCM vs. FTM (Fig. 6B, right panel).

**Figure 6.**
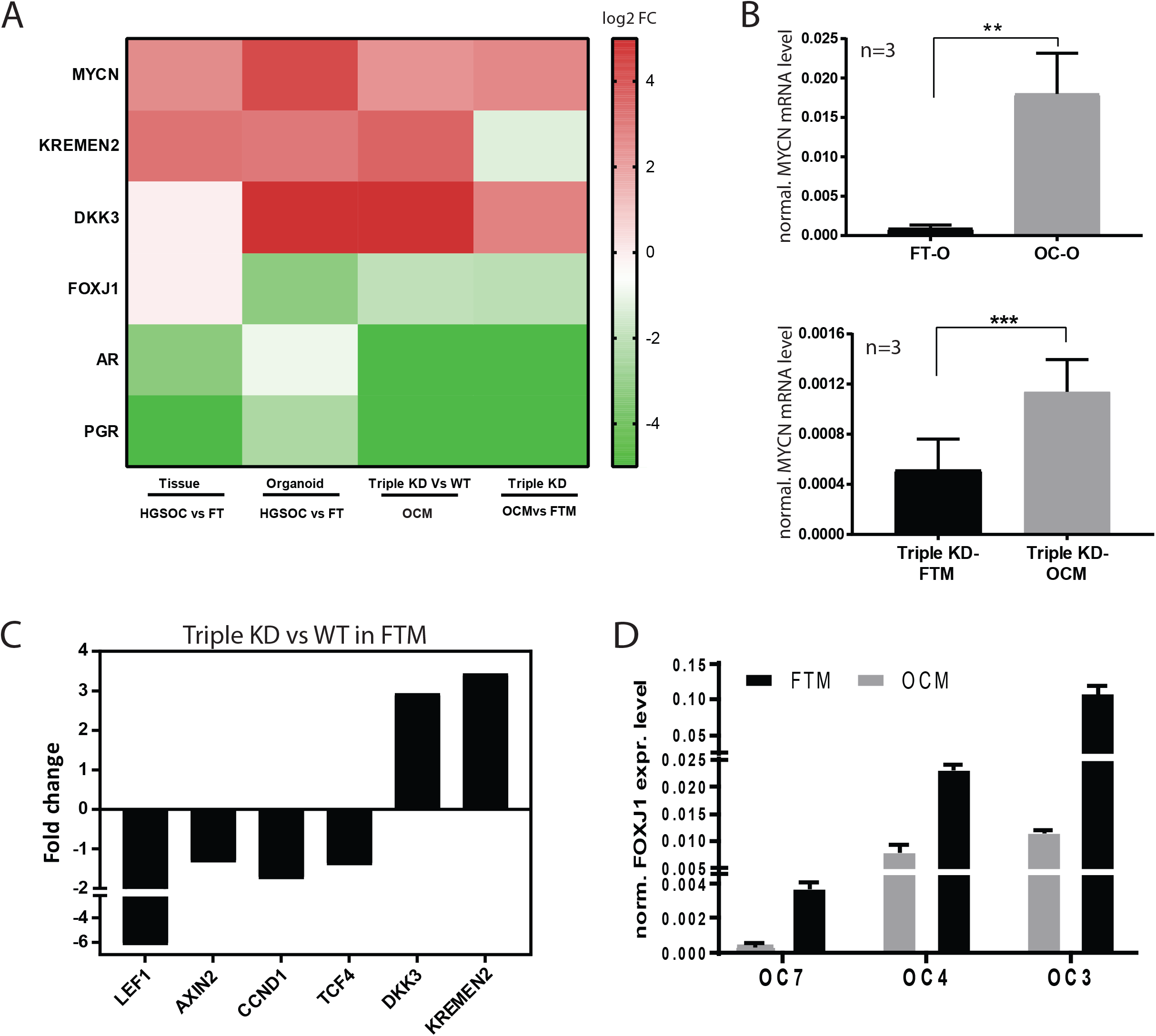
Unifying induction of MYCN and Wnt inhibitors in HGSOC and KD samples. A) Comparative analysis of microarray data revealed consistent up-regulation of MYCN, the canonical Wnt inhibitors KREMEN2 and DKK3 as well as differentiation marker FOXJ1, AR and PGR in HGSOC cancer organoids as well as triple KD FT organoids. Differential expression was determined either by single-color microarray for the cancer samples (8 replicates) or dual-color microarray for the knockdowns (2 replicates) and is significant for all genes with p < 0.05. B) Transcriptional induction of MYCN is confirmed by qRT-PCR for three patients between cancer organoids and normal FT organoids as well as triple KD organoids grown in OCM vs. FTM conditions. Depicted is the mean ±SEM of the normalized MYCN mRNA expression level of three biological replicates (n=3). ** p < 0.01, *** p < 0.001, two-sided Student’s t-test. C) Microarray data revealed that expression levels of Wnt target genes are downregulated in triple KD compared to WT control organoids. Differential expression determined by dual-color microarray for 2 biological replicates was significant for all genes with p ≤ 0.005. D) FOXJ1 expression was upregulated in ovarian cancer organoids in FTM compared to OCM as shown by qRT-PCR analysis of 3 different patient samples. Data represent the mean ±SEM of technical triplicates.

In line with our finding that suppression of canonical Wnt signaling supports the growth of cancer organoids, the gene expression data revealed enhanced expression of the Wnt inhibitors KREMEN2 and DKK3 in both triple KD and OC organoids (Fig. 6A). While KREMEN2 regulation appears to be driven by changes in the genetic background, DKK3 expression seems to be induced by Wnt-free medium. Therefore, it appears that the altered paracrine signaling environment further promotes inhibition of canonical Wnt signaling. Interestingly, downregulation of p53/PTEN/RB in FT organoids caused a decrease in Wnt target gene expression even in the presence of Wnt3a and RSPO1 (Fig.6C), while culture in Wnt-free medium leads to a further strong reduction of LEF1 and TCF transcripts down to the detection limit of the microarray (see Supplementary Table 6). The expression of Wnt inhibitors and the reduced Wnt signaling induced by tumor suppressor KD are in agreement with our hypothesis that Wnt pathway suppression is required for HGSOC growth. Interestingly, HGSOC organoids expressed increased levels of differentiation markers when exposed to Wnt3a/RSPO1. In line with the microarray results and data from the triple KD organoids, RT-PCR confirmed increased expression of the ciliogenesis factor FOXJ1 in cancer organoids cultured in the presence of Wnt3a/RSPO1 (Fig. 6D). Based on this, we postulate a two-step process of HGSOC development in which changes in the niche environment are a necessary step that drives dedifferentiation and maintains stemness in cells bearing tumorigenic mutations.

## Discussion

Our understanding of the cellular mechanisms that underlie the development of HGSOC in the FT and its spread to the ovaries and beyond remains rudimentary and represents a major obstacle to advances in diagnosis and therapy. Although some properties of cancer stem cells in HGSOC have been described (Choi et al., 2015; Lupia and Cavallaro, 2017; McLean et al., 2011), there is no knowledge about the intrinsic mechanisms that drive tumor growth and mediate recurrent disease. As patients frequently receive neoadjuvant chemotherapy prior to debulking surgery, such tumor samples are likely to already show secondary biological alterations. This study thus provides systematic insight into the biology of HGSOC organoids derived directly from debulking surgery samples, which reflect the biology of the initial cancer tissue.

In contrast to the healthy FT epithelium (Kessler et al., 2015), we find that HGSOC cancer organoids require a low Wnt signaling environment but an active BMP signaling axis. Wnt independence was previously observed in other cancers such as pancreas and colon (Fujii et al., 2016; Seino et al., 2018) and positively correlates with the stage of metastatic disease. However, in contrast to organoids from other cancers, e.g. metastatic pancreatic cancer, which do not require Wnt but do grow in Wnt/Noggin/Rspo medium, exogenous supplementation of Wnt3a actually prevented formation and growth of all HGSOC cancer organoids. Recently, Kopper et al (2019) also reported the use of Wnt-free medium for the generation of HGSOC organoids but attributed negative effect of Wnt to the potential contaminating presence of serum without testing the hypothesis. Here we clearly show that lack of exogenous Wnt signaling is in fact a requirement for maintaining stemness and preventing differentiation in HGSOC and that pro-cancerous mutations by themselves are therefore likely not sufficient to drive transformation of healthy epithelium. Thus, it is tempting to speculate that the overall environment in the FT epithelium (with presumably high Wnt signaling) represses cancer outgrowth and promotes escape of (pre)-malignant cells to more distant sites, like the surface of the ovaries and peritoneum. Interestingly, even formation of the two organoid cultures that benefited from the presence of RSPO1 (Figure 1) could be efficiently prevented by addition of Wnt3a and noggin, suggesting high sensitivity of the HGSOC organoids to elevated Wnt signaling. Nevertheless, these cases show that the heterogeneity of this disease warrants routine testing of two different media during establishment of fresh cultures: OCM and OCM + RSPO1 and FGF.

Our analysis of HGSOC and KD organoids clearly demonstrates that a high Wnt environment leads to the downregulation of stemness genes and the upregulation of differentiation genes. Mouse lineage tracing data previously showed that active Wnt signaling is required both for renewal of Pax8 secretory cells and their differentiation to ciliated cells (Ghosh et al., 2017). Our data strongly suggest that the regulation of these processes becomes separated during carcinogenesis. While Wnt stimulation still induces differentiation and ciliogenesis in cancer organoids, it inhibits stemness and expansion capacity. The same effect is observed in triple KD organoids, indicating that depletion of key tumor driver proteins is sufficient to induce these alterations in stem cell regulation. Yet, despite the change in growth requirements, the prominent occurrence of nuclear atypia, increase of DNA damage and changes in epithelial organization, triple KD organoids maintained key elements of epithelial polarity. Therefore, it appears they have not completed the process of transformation and that additional changes are needed for malignancy to develop. This is also in line with the observation that growth rescue in OCM medium was incomplete. Unlike the stable long-term expansion of HGSOC organoids, which were routinely cultured for over 1 year without changes in growth dynamics, the lifespan of KD organoids in OCM was limited to 7-8 months.

Our data illustrate the critical importance of the model system when studying processes of cell transformation *in vitro*. Several previous studies on the putative role of the FT as the tissue of origin of HGSOC showed the transformation potential of human FT epithelial cells in 2D cell culture by overexpression of different oncogenes (h-RAS, c-MYC) followed by xenograft transplantation (Jazaeri et al., 2011; Karst et al., 2011). We show that robust regulatory mechanisms in the intact epithelium prevent the breakdown of the epithelial organization despite functional inactivation of the key HGSOC drivers p53, PTEN, and RB. Breakdown of apicobasal polarity is an important step in the emergence of many cancers, as it precedes EMT and thus facilitates cancer progression (Huber et al., 2005; Ozdamar et al., 2005). While our microarray data revealed important similarities in gene expression profile between KD FT organoids and mature cancer organoids, further studies are needed to elucidate the exact mechanism of cancer stem cell maintenance in HGSOC. MYCN, which in the absence of Wnt was ubiquitously upregulated in all OC and KD FT organoids in our study, was previously implicated as a driver of stemness in aggressive glioblastoma (Yang et al., 2017). The high degree of consistency in MYCN regulation in KD FT and HGSOC organoids from independent donor samples warrants further studies to investigate its potential role as a stemness marker in HGSOC.

Despite immense efforts to establish new *in vitro* models of HGSOC (Hill et al., 2018), a successful model for long-term expansion of solid tumor deposits has been missing until now. This is likely due to the altered stem cell niche requirements we describe here. Our preliminary tests with carboplatin showed individual differences in drug response of organoids from different patients, suggesting that the model is suitable for personalized therapeutic strategies. However, to define to which extent organoids are predictive of *in vivo* patient responses, more comprehensive studies in correlation with long-term clinical outcome are needed. It is possible that including other cell types in the organoid model, such as stromal and immune cells, could further advance its capacity to generate data which are predictive of clinical responses in patients (Neal et al., 2018).

Overall this study provides important insight into fundamental biological processes of HGSOC development. We show that functional inactivation of key tumor drivers fails to induce a direct growth advantage of the altered cells in the absence of appropriate changes in the stem cell niche environment. The existence of such a two-component mechanism opens up important new questions for understanding the etiology of HGSOC – in particular which cellular and physiological mechanisms in the tissue surrounding the FT epithelium are responsible for the critical changes in paracrine signaling that favour the outgrowth of mutated cells. In this context, the regulatory role of the neighboring ovary could further be of pivotal relevance, particularly in light of epidemiological data showing a strong correlation between inflammatory processes associated with ovulation and the risk for HGSOC development.

## Acknowledgements

We would like to thank Susan Jackisch, Ina Wagner, Jörg Angermann and Oliver Thieck for technical support, Dr Gabriela Vallejo Flores, Toralf Kaiser and Jenny Kirsch for technical help with FACS experiments, Diane Schad for expert help with generating graphics and Dr. Rike Zietlow for editing the manuscript.

## Author contributions

M.K. and T.F.M. conceived the project; M.K. and K.H. designed experiments, which were conducted by, K.H., M.K and S.T.; H.K. designed and planned targeted sequencing, performed by T.Z. H.B. and T.Z. analyzed the targeted sequencing data. H.B. and H.-J.M. analyzed microarray data. S.D.E and E.T performed pathology analysis of the tissue samples. M.M., selected patients and provided human FT samples J.S. and E.B. selected the patients, provided material, and clinical data of primary HGSOC patients. T.F.M. supported the project financially; K.H, M.K. and T.F.M. wrote the manuscript, M.K. and T.F.M. supervised the project.

## Declaration of Interests

The authors declare no competing interests.

## Materials and Methods

### Fallopian tube and ovarian cancer primary patient material

Approval for the preparation and experimental usage of the primary material was given by the Ethics Commission of the Charité, Berlin (EA1/002/07) and informed consent was obtained from every patient.

Human fallopian tube samples were provided by the Auguste-Viktoria Klinikum Berlin, Department of Gynecology and Obstetrics after standard surgical procedures for benign gynecological disease. Only anatomically normal fallopian tubes were used. The tubes were transported and dissected within 2 to 3 h of removal. Of each FT sample one piece was saved for RNA/DNA preparation. After washing with DPBS, FT tissue was incubated in collagenase I (Sigma) for 45-60min at 37°C for enzymatic detachment of epithelial progenitors. Subsequently, the mucosal cells were scraped off the muscularis with a scalpel and pelleted by centrifugation (7min, 300xg). Cells were seeded in 2D culture (in ADF+++ with Pen/Strep, hEGF and ROCK inhibitor) before transfer into Matrigel™ for organoid formation.

Ovarian cancer tissue was received from the Department of Gynecology, Charité University Hospital, Campus Virchow Clinic. Cancer samples, surgically removed from patients with a preliminary diagnosis of HGSOC were retrieved from tumors of specified sites within the abdominal cavity. The tumor material was transported and processed within few hours after surgery and each sample was divided into 3 pieces for cell preparation, tissue fixation and RNA/DNA isolation. For isolation of HGSOC progenitors, the tissue pieces were washed with DPBS, minced with a scalpel into very small pieces and incubated in a 1:1 mixture of collagenase I and II (Sigma) for around 60 min at 37°C on a shaker. Afterwards, the enzyme-tissue mixture was vortexed to further separate the cells. Next, the cell suspension was centrifuged (5 min, 300 x g, 4°C), resuspended in ADF+++ supplemented with Pen/Strep, hEGF and ROCK inhibitor and seeded in a cell culture vessel or directly in Matrigel™.

### Organoid culture of FT epithelial cells (FTECs)

The organoids were cultured as described in Kessler et al (2015). In brief, to initiate organoid growth approximately 30,000 cells obtained from 2D culture, were seeded in 50 µL Matrigel™ and overlaid with a growth factor cocktail stimulating different paracrine pathways including EGF and Wnt signalling. The composition for normal FT epithelial cell medium (FTM) was as follows: ADF, 25% conditioned Wnt3A-medium and 25 % conditioned RSPO1 medium, supplemented with 12 mM HEPES, 1% GlutaMAX™, 2% B27, 1% N2, human EGF, human noggin, human FGF-10, 1 mM nicotinamide, 9 µM ROCK inhibitor (Y-27632) and 0.5 µM TGF-β RI Kinase Inhibitor IV (SB431542). Medium was changed every 3-4 days and organoids were expanded every 2-3 weeks at a rate of 1:2 to 1:3. To do so, organoids were released from Matrigel™, washed with ADF++, taken up in pre-warmed TrypLE™ and incubated for 7-10 min at 37 °C for enzymatic digestion. Subsequently, organoids were vortexed, washed with ADF++ and pelleted. Finally, the shredded organoids were resuspended in fresh Matrigel™ and seeded into pre-warmed cell culture plates before FTM was added to the organoids.

### Organoid culture of ovarian cancer cells

After initial isolation from the tumor tissue, ovarian cancer cells were seeded in Matrigel™ at a concentration of approx. 30,000 cells/50 µl. Because cancer cells did not form organoids or were difficult to expand in FTM, different media compositions were tested to support *in vitro* growth. The basic medium for the ovarian cancer organoids always contained the following supplements and growth factors: ADF supplemented with 12 mM HEPES, 1% GlutaMAX™, 2% B27, 1% N2, human EGF, 1 mM nicotinamide, 9 µM ROCK inhibitor (Y-27632) and 0.5 µM TGF-β RI Kinase Inhibitor IV (SB431542). Upon proper organoid maturation and growth under specified conditions they were split at a ratio of 1:3 every 1-2 weeks. To expand the OC organoids they were released from Matrigel™ using ice-cold ADF++, centrifuged and resuspended in pre-warmed TrypLE™. Enzymatic digestion was carried out for 7 min at 37 °C. Afterwards, the organoid suspension was vortexed, mixed with cold ADF++ and centrifuged (300xg, 5 min). Next, the supernatant was removed, the cell pellet resuspended in Matrigel™ and seeded into pre-warmed cell culture plates. After ∼20 min the Matrigel™ was overlaid with the respective medium. The medium was changed twice per week and the organoids kept in a humidified incubator at 37 °C and 5% CO_2_.

| Medium/Supplement | Provider (Cat. no.) | Final conc. |
| --- | --- | --- |
| **Advanced DMEM/F12 (ADF)** | Gibco (12634) |  |
| **B27 supplement** | Gibco (17504044) | 1x |
| **BMP2, human** | Gibco (PHC7146) | 10 ng/ml |
| **EGF, human** | Gibco (PHG0311) | 10 ng/ml |
| **Fetal calf serum (FCS)** | Biochrom (S0115) | 5 or 10% |
| **FGF-10, human** | Peprotech (100-26-B) | 100 ng/ml |
| **GlutaMax** | Gibco (35050-038) | 1x |
| **HEPES** | Gibco (15630-056) | 12 mM |
| **N2 supplement** | Gibco (17502048) | 1x |
| **Nicotinamide** | Sigma-Aldrich (N0636) | 10 mM |
| **Noggin, human** | Peprotech (120-10C) | 100 ng/ml |
| **Nutlin3a** | Sigma (N6287) | 5-10 μM |
| **Penicillin/Streptomycin** | Gibco (15140-122) | 100 U/ml<br>10 mg/ml |
| **Puromycin** | Gibco (A11138-03) | 0.5 μg/ml |
| **Y-27632 dihydrochloride monohydrate (ROCK inhibitor)** | Chemdea (CD0141) | 9 μM |
| **Rspo-1** | In-house production<br>Cell line: 293T HA Rspo1-Fc | 10-25% (v/v) |
| **SB431542 (TGF-β inhibitor)** | Calbiochem (616454) | 0.5 μM |
| **Wnt3a** | In-house production (Willert et al. 2003)<br>Cell line: L Wnt-3a (ATCC® CRL-2647™) | 10-25% (v/v) |

### Organoid stocks

Organoid stocks were prepared by releasing organoids from ∼1 week-old confluent cultures in cold ADF++, pelleting by centrifugation and resuspension in the freezing medium CryoSFM. After transfer of the organoid suspension into cryotubes and a freezing container, stocks were kept at least 24 h at - 80 °C before transfer into liquid nitrogen for long-term storage.

### Cloning of shRNAs and virus production

To obtain stable knockdowns of PTEN and RB (in-house-designed) shRNA sequences targeting specific regions within the respective mRNA were cloned into the lentiviral vector pLVTHM (Addgene#12247) carrying mCherry or GFP as mammalian selection marker. Cloning included the following steps: 1. *Clal*/*Mlul* (NEB) restriction of pLVTHM-GFP or –mCherry vector, 2. Annealing of sense and antisense oligos (target sequence with specific overhangs), 3. Ligation of the linearized vector and annealed oligos (Quick ligation kit, NEB) and 4. Transformation into *E.coli*. The retroviral vector with the shRNA against p53 was provided by Alexander Loewer.

**Replication-deficient retro- and lentiviral particles were produced** by transient transfection of 293T cells using FuGENE® 6 reagent (Promega). In brief, specific amounts of viral target and packaging plasmids were mixed with Opti-MEM™ (Thermo Scientific). Next, a mixture of Opti-MEM™ and FuGENE® 6 was added to the plasmids and incubated for 20-30 min at RT. The formed liposomes were added dropwise to 293T cells (∼75% confluence) in 1xDMEM (supplemented with 2 mM L-glutamine, 1mM Na-pyruvate and 10% FCS). The next day (∼12 h), the transfection medium was aspirated and fresh medium added. Around 36-48h post-transfection the viral supernatant was collected, filtered (0.45 µm) and concentrated with Lenti-X™ concentrator (Clontech). The viral pellet was resuspended in ADF medium to achieve 10x concentrated virus. Aliquots were stored at −80°C.

### Lentiviral vectors

psPAX2 (packaging plasmid; Addgene#12260), pMD2.G (VSV-G envelope plasmid; Addgene#12259), pLVTHM-GFP (Addgene#12247), pLVTHM-GFP/mCherry-Luci (shRNA against luciferase as control vector; target sequence (5’-3’): *AACTTACGCTGAGTACTTCGA*), pLVTHM-mCherry-PTEN (pLVTHM-mCherry with shRNA against PTEN; target sequence (5’-3’): *AGGCGCTATGTGTATTATTAT*), pLVTHM-GFP-Rb (pLVTHM-GFP with shRNA against Rb; target sequence (5’-3’): *GGTTGTAATGGCCACATATAG*)

Retroviral vectors were a gift by Alexander Loewer and plasmids are described in Brummelkamp et al. (2002) (retroviral packaging plasmid, retroviral envelope plasmid, pSUPER.retro (Addgene#3926), pSUPER.retro.p53 (pSUPER.retro with shRNA against p53; target sequence (5’-3’): *GACTCCAGTGGTAATCTA*C)

### Genetic manipulation of primary cells

To genetically modify primary FT epithelial cells, they were treated with retro-and lentiviruses in 2D culture. Freshly isolated FECs of an early passage (p0/p1) were transduced when reaching around 50-70% confluence with 1x concentrated virus diluted in ADF+++ and supplemented with hEGF and 5 µg/µl polybrene (Sigma). The cells were incubated for 12-18 h with the viral suspension. The next day, the supernatant was removed and fresh 2D medium added to the cells. When the cultures reached 80-90% confluence they were split with TrypLE™ and seeded back into 2D at a ratio of 1:2 to 1:3. In the next passage the second virus was added in the same manner. Cells were sequentially transduced and expanded until they reached max. passage 3.

To retrieve successfully transduced cells carrying a fluorescent selection marker, FACS was performed. For this purpose, the primary cells were detached using TrypLE™ (∼10 min), pelleted by centrifugation (5min, 300xg), resuspended in FACS buffer (1xDPBS with 1% FCS, 1% HEPES and 3% ROCK inhibitor) and passed through a 35 µm cell strainer into polystyrene tubes.

Propidium iodide (Sigma) was added shortly before FACS to exclude the dead cells. Sorting was performed by the Flow cytometry core facility (DRFZ, Charitéplatz 1, 10117 Berlin) using the FACSAria II (BD). The GFP and mCherry single or double-positive cells were collected in polystyrene tubes supplied with 2D medium. Collected cells were centrifuged directly after FACS and seeded in 3D culture with around 1.5 × 10^5^ cells/ 50 µl Matrigel™. Cells transduced with the retroviral vector were selected by adding 0.5 µg/ml puromycin (Gibco) for 10 days to the respective cell culture.

### Single cell preparation and flow cytometry

To prepare single cells, organoids were released from Matrigel™ using cold PBS, pelleted by centrifugation (5 min, 300xg), resuspended in TrypLE and incubated for 15-20 min at 37 °C. After enzymatic treatment and vortexing, the organoid fragments were further mechanically disrupted by passing 3-4x through a needle (26G). Next, the cells were taken up in ADF and passed through a 40 µm filter. The single-cell suspension was pelleted and washed by addition of 1% BSA in PBS. The cells were stained in 1xPBS with 1-2% BSA. Staining was performed for 10-30 min at 4 °C in the dark. Finally, the cells were washed and resuspended in 1xPBS. Flow cytometric analysis was performed using a FACSCanto™ II (BD) or LSRFortessa™ (BD) flow cytometer and the FlowJo vX.0.6 software.

### Cell viability assay

To quantify the number of viable cells within one organoid well, we applied the CellTiter-Glo® 3D Cell Viability Assay (Promega # G9681). The assay was carried out according to the manufacturers’ protocol and the luminescence was measured within 30 min after the start of the reaction using black 96-well plates (Costar®, Corning) and a standard plate reader.

### Immunohistochemistry of HGSOC tissue

Paraffin-embedded tumor samples in 5 µm sections was stained with the mouse monoclonal antibodies EpCam (clone VU-1D9, Thermoscientific, Thermo Fischer Scientific, Waltham, Massachusetts, USA) at 1:100, for p16 (clone 16P04, NeoMarkers, Fremont, Calfornia, USA) at 1:600, for p53 (clone DO-7, Dako, Carpinteria, California, USA) at 1:50 dilution and for Pax8 (clone MRQ-50, Cell Marque by Sigma-Aldrich, Rocklin, California, USA) at suppliers instructions on the Ventana Benchmark XT Autostainer instrument (Ventana Medical Systems, Inc., Tucson, AZ, USA). 3, 3’-diaminobenzidine peroxide substrate (DAB^+^) of the “ultraView Universal DAB detection kit” (Ventana) was used as a chromogen. Signals were strong and clearly discernable and furthermore well-established in our diagnostic routine laboratory.

### Immunofluorescence staining

For fixation organoids were released from Matrigel and incubated in 3.7% PFA for ∼1 h at RT, while tissue pieces were incubated in PFA for ∼24 h at RT. Organoids and tissues were then dehydrated and paraffinized. For dehydration organoids were passed manually through an alcohol dilution series: 60% EtOH (20 min, RT), 75% EtOH (20 min, RT), 90% EtOH (20 min, RT), abs. EtOH (20 min, RT), 100% isopropanol (20 min, RT) and 2x 100% acetone (20 min, RT). Tissue pieces were transferred to the Shandon Citadell 1000 rondell for automatic dehydration. After dehydration, organoids and tissue pieces were embedded in paraffin, sectioned at 5 µm and collected onto microscope slides.

Prior to staining, the formaldehyde-fixed paraffin-embedded (FFPE) organoids and tissues were deparaffinized by applying a series of decreasing alcohol concentrations: xylene (2x 10 min), 100% ethanol (2x 2 min), 90% ethanol (1x 2 min), 70% ethanol (1x 2 min), 50% ethanol (1x 2 min) and H_2_O (2x 2 min). For antigen retrieval the slides were incubated for 30 min at 98 °C in 1x Target retrieval solution (Dako). Sections were stained by adding primary antibodies diluted in immunofluorescence buffer (IFB: 1% BSA, 2% FCS and 0.1% Tween-20 in 1xPBS) and depending on the antibody incubated for 90 min at RT or overnight at 4°C in a humidified chamber. After washing 5x 5 min with 1x PBS supplemented with 0.05% Tween-20, the respective secondary antibodies diluted in IFB together with the nucleic acid dye DR or Hoechst were added to each section and incubated for 60 min at RT. Subsequently, the slides were washed and sections mounted on coverslip using Mowiol.

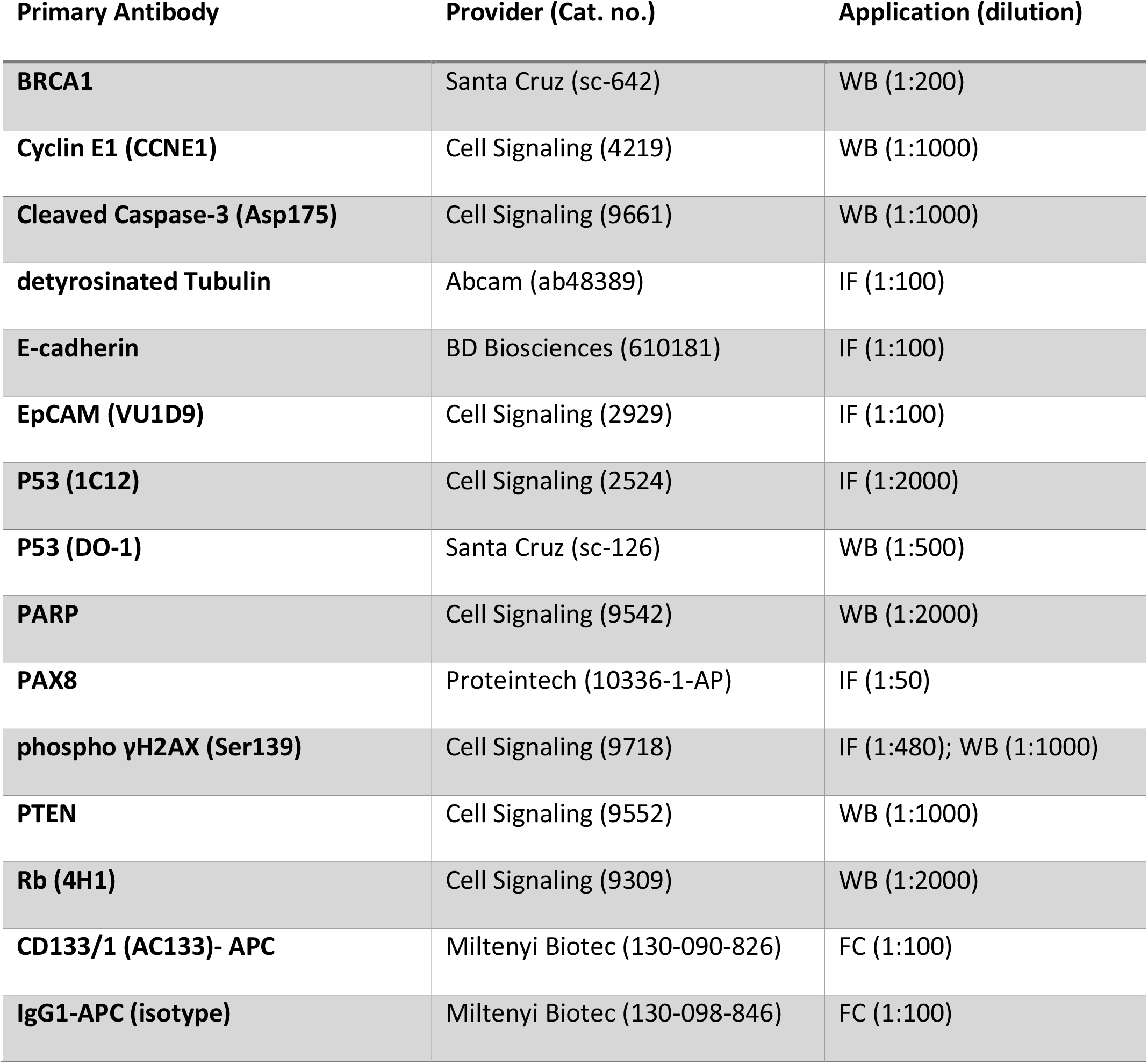

### H&E-Staining

Before staining with haematoxylin and eosin (H&E), organoid slices were deparaffinized as described above. After deparaffinization, the sections were covered completely with Mayer’s haematoxylin solution (Roth) and incubated for 15 min at RT. The slides were rinsed with ddH2O and tap water. Subsequently, the sections were covered with Eosin Y solution (1% aqueous, Roth) and incubated for 10 min at RT. Finally, slides were rinsed for 1 min with ddH2O and embedded using Roti®-Histokitt.

### Quantitative Reverse-Transcription PCR (qRT-PCR)

The AB Power SYBR® Green RNA-to-CT™ 1-Step Kit (Thermo Fisher) was applied to perform reverse transcription and quantitative PCR in one step. RNA was added in a volume of 10 µl at a concentration between 2-10 ng/µl. qRT-PCR was performed using the StepOnePlus™ Real-Time PCR System (Applied Biosystems) with the following program: 1. 30 min, 48 °C; 2. 10 min, 95 °C; 3. 15 sec, 95 °C and 4. 1 min, 60 °C. Steps 3 and 4 were repeated 40 times. For each primer pair and RNA-sample the reaction was done in triplicate. The amplification plots obtained from the qRT-PCR were analyzed with the StepOne™ Software (version 2.3; Thermo Fisher). The expression levels were relatively quantified by calculating ΔΔCt. The expression levels of the target genes were always normalized to the expression of the housekeeping gene glyceraldehyde-3-phosphate dehydrogenase (GAPDH).

Primer sequences (5’-3’): GAPDH_for – GGTATCGTGGAAGGACTCATGAC; GAPDH_rev – ATGCCAGTGAGCTTCCCGTTCAG; Ki67_for – AAGCCCTCCAGCTCCTAGTC; Ki67_rev – TCCGAAGCACCACTTCTTCT; FOXJ1_for – GGAGGGGACGTAAATCCCTA; FOXJ1_rev – GGTCCCAGTAGTTCCAGCAA; TP53_for – CCTCCTCAGCATCTTATCCGA; TP53_rev – TGGTACAGTCAGAGCCAACCTC; PTEN_for – CGACGGGAAGACAAGTTCAT; PTEN_rev – AGGTTTCCTCTGGTCCTGGT; RB_for – GGAAGCAACCCTCCTAAACC; RB_rev – TTTCTGCTTTTGCATTCGTG; AXIN2_for – AGATCCAGTCGGTGATGGAG; AXIN2_rev – CTTCATTCAAGGTGGGGAGA; MYCN_for – CTTCGGTCCAGCTTTCTCAC, MYCN_rev - GTCCGAGCGTGTTCAATTTT

### Western blotting

For the purpose of sample collection organoids were washed with DPBS, pelleted and lysed by addition of 1x Laemmli buffer and heating for 10 min at 95 °C. After the proteins were separated according to their molecular weight by performing SDS-PAGE, they were transferred from the gel to a PVDF membrane using a Mini Trans-Blot® Electrophoretic Transfer Cell (Bio-Rad) and applying 250 mA, 400 V and 4 °C under constant stirring for 2.5 h. After blocking in a mixture of 5% milk and BSA the membrane was incubated at 4 °C overnight with the primary antibody. The next day the membrane was washed 3x 10 min in TBST. Subsequently, the membrane was incubated for 1 h at RT with the respective conjugated secondary antibody. After washing, the membrane was covered with chemiluminescence reagents. Using Hyperfilm (Amersham Biosciences) and a developer machine, the proteins were visualized. The housekeeping gene β-actin was used as the internal control for normalization of protein loading.

### DNA/RNA isolation

DNA was isolated using either the AllPrep DNA/RNA Mini Kit (QIAGEN) for tissue and organoids. DNA purifications were quantified by measuring optical density at 260 nm wavelengths.

RNA from cells of 3D cultures was isolated using the RNeasy Mini Kit (QIAGEN) or the AllPrep DNA/RNA Mini Kit (QIAGEN). RNA from tissue samples was isolated using the AllPrep DNA/RNA Mini Kit (QIAGEN). After pelleting the organoids, they were lysed in the respective sample buffer and RNA purification was performed according to the manufacturer’s protocols. RNA concentration and purity were measured with a NanoDrop® ND-1000 Spectrophotometer.

For microarray analysis total RNA was isolated with TRIzol (Life Technologies) according the supplier’s protocol using glycogen as carrier. Quality control and quantification of total RNA was carried out using the NanoDrop 1000 UV-Vis spectrophotometer (Kisker) as well as the Agilent 2100 Bioanalyzer with a RNA Nano 6000 microfluidics kit (Agilent Technologies).

### Drug testing

To test the response of patient-derived HGSOC organoids to the chemotherapeutic drug carboplatin (Merck), organoids were dissociated into single cells by using enzymatic (TrypLE, 15 min, 37 °C) and mechanical (vortex and passing through 26G needle) disruption. Subsequently, cells were counted and seeded at 15,000 cells/ 25µl Matrigel™ into a 48-well plate. After maturation of organoids (∼7-10 days in culture) treatment with carboplatin with a concentration range between 0-100 µg/ml was started. At one week post-treatment, cell viability was determined as described above.

### Microarrays (Dua-color and Single-color)

Microarray experiments were performed as dual-color or single-color hybridizations on either Agilent Whole Human Genome 4×44K microarrays (Design ID 014850) or 8×60K human custom (Agilent-048908) microarrays comprising identical features for coding genes. Color-swap dye-reversal hybridizations were performed in order to compensate for dye specific effects and to ensure statistically relevant data when using small sample sizes. RNA labeling was done either with a two-color Quick Amp Labeling Kit (4×44K arrays) or with a one-color Low Input Quick Amp Kit (8×60K arrays) according the supplier’s recommendations (Agilent Technologies).

In brief, mRNA was reverse transcribed and amplified using an oligo-dT-T7 promoter primer and the T7 RNA polymerase. The resulting cRNA was labeled with only Cyanine 3-CTP (single-color) or with Cyanine 3-CTP and Cyanine 5-CTP (dual-color). After precipitation, purification, and quantification, 1 μg (4×44K arrays) or 300 ng (8×60K arrays) of each labeled cRNA was fragmented and hybridized to whole-genome multipack microarrays according to the manufacturer’s protocol (Agilent Technologies). Scanning of microarrays was performed with 5 μm resolution and XDR extended range (4×44K arrays) or 3 µm resolution (8×60K arrays) using a G2565CA high-resolution laser microarray scanner (Agilent Technologies). Microarray image data were analyzed and extracted with the Image Analysis/Feature Extraction software G2567AA v. A.11.5.1.1 (Agilent Technologies) using default settings and either the GE2_1100_Jul11 (dual-color) or the GE1_1105_Oct12 (single-color) extraction protocol.

For analysis of dual-color microarrays the extracted MAGE-ML files were processed with the Rosetta Resolver, Build 7.2.2 SP1.31 (Rosetta Biosoftware). Ratio profiles comprising single hybridizations were combined in an error-weighted fashion to create ratio experiments. A 1.5-fold change expression cut-off for ratio experiments was applied together with anti-correlation of ratio profiles, rendering a highly significant, robust and reproducible microarray analysis (p-value < 0.01). Additionally, raw data txt files were analyzed with R packages from the Bioconductor repository.

The extracted single-color raw data files were background corrected, quantile normalized and further analyzed for differential gene expression using R 3.4 (Ritchie et al., 2015) (Supplementary Information Table 2 and 3). Microarray gene expression comparisons between groups and the associated BioConductor package LIMMA were performed using unpaired tests for all human comparisons (R Core Team, 2013).

### Capture of the targeted disease-related genome and Next-Generation Sequencing

A SureSelectXT Automation Custom Capture Library (Agilent) target enrichment panel was designed. The enrichment panel comprised all coding exons of 121 genes associated with ovarian cancer (Supplementary Table 3). Capture was performed according to the manufacturer’s instructions using an NGS Workstation Option B (Agilent) for automated library preparation starting with 3 µg DNA per sample. Sequencing was performed on an Illumina HiSeq 2500 system generating 2×100bp paired end reads with a target coverage of >200-fold per sample. Sequence reads were mapped to the haploid human reference genome (hg19) using BWA. Single nucleotide variants (SNVs) and short insertions and deletions (indels) were called using FreeBayes v1.1. (Garrison and Marth, 2012).

Variants called by FreeBayes were filtered for quality (QUAL > 10, coverage > 50) and annotated by SnpEff v4.3k (Cingolani et al., 2012) and Annovar (Wang et al., 2010). For each variant the effect with the highest impact as defined by SnpEff was selected. Variants were flagged as rare if they showed less than 1% population frequency in the 1000 genome (Auton et al., 2015) and ESP6500 [Exome Variant Server, NHLBI GO Exome Sequencing Project (ESP), Seattle, URL: http://evs.gs.washington.edu/EVS/; accessed 2014-12] data sets. Predictions of amino acid exchange effects on protein function from MetaSVM, MetaLR and M-CAP as provided by Annovar were used to assess loss of function.

### Code Availability

All code used for generating analyses used in this publication is available from https://github.com/MPIIB-Department-TFMeyer/Hoffmann_et_al_Ovarian_Cancer_Organoids.

