## Supplementary Information for "Preservation of stemness in high-grade serous ovarian cancer organoids requires low Wnt environment"

#### Supplementary Figure Legends

**Supplementary Figure 1. Pathology of HGSOC tumor samples and drug response of HGSOC organoids.** A) Diagnosis of ovarian cancer in tissue fragments used for organoids generation were confirmed by experienced pathologists to be of HGSOC phenotype, being marked by strong EpCAM, Pax8 and p16 expression. Depicted are three examples of the pathology stainings. B) Three HGSOC organoid lines were tested for their response to the drug carboplatin showing slightly different responses as determined by cell viability assay. Data represent mean ±SD of technical triplicates.

**Supplementary Figure 2. Expression profile of HGSOC organoids.** A) Representative western blot indicating loss of BRCA1 protein expression in sample OC4 but WT level expression in OC7. B) Nuclear p53 expression in different cancer organoids was confirmed by IF stainings. C) Spearman correlation map based on single-color microarray data indicates stronger similarity in expression patterns between the healthy organoids vs tumor organoids**.** D) Protein expression profiling confirmed frequent CCNE1 overexpression in established HGSOC organoid lines.

**Supplementary Figure 3. Effects of p53, PTEN and Rb knockdown in FT organoids.** A) Gating strategy to sort GFP/mCherry double-positive single cells derived from transduced FT organoids. B) Activation of the AKT pathway as indicated by increased levels of phospho Akt protein proves functional PTEN knockdown. C) P53 KD organoids are resistant to Nutlin3A treatment, confirming functional disruption of p53 signaling. D) Summary table and representative phase contrast images of the 7 triple KD organoid lines that were generated from different donors, all of which showed a premature growth arrest.

**Supplementary Figure 4. Confirmation of DNA damage and apoptosis as well as activation of Wnt target genes.** A) Growth curves of a second biological replicate showing loss of growth capacity and premature growth arrest of KD organoids compared to the controls when grown in FTM. B) Phase contrast images of 3 engineered organoid lines (p53/PTEN/RB knockdown) differentially treated with FTM and OCM confirm differences in organoid formation and long-term growth. Scale bar: 500 µm. C) Western blot from donor FT268 as independent biological replicate of the effects shown in Fig. 4C. Increase in DNA damage (γH2AX, PARP1 and cleaved CC3) in triple KD organoids is not significantly affected by the medium change. D) Wnt target genes were significantly downregulated upon change from FTM to OCM as revealed by microarray analysis. Differential expression determined by dual-color microarray for 2 biological replicates was significant for all genes with $p < 0.05$.

**Supplementary Figure 5. HGSOC organoids respond to exogenous Wnt signals with changes in gene expression and growth pattern.** A) The percentage of CD133+ cells was determined by flow cytometry in knockdown and control organoids. B) Gating strategy for CD133+ cells in HGSOC organoids. C) Canonical Wnt target gene AXIN2 is significantly upregulated in HGSOC organoids treated with Wnt3a/Rspo1 conditioned medium as confirmed by qRT-PCR. AXIN2 expression levels were normalized to GAPDH and calculated as mean ±SEM from technical triplicates.

Supplementary Figure 1. Pathology of HGSOC tumor samples and drug response of HGSOC organoids.

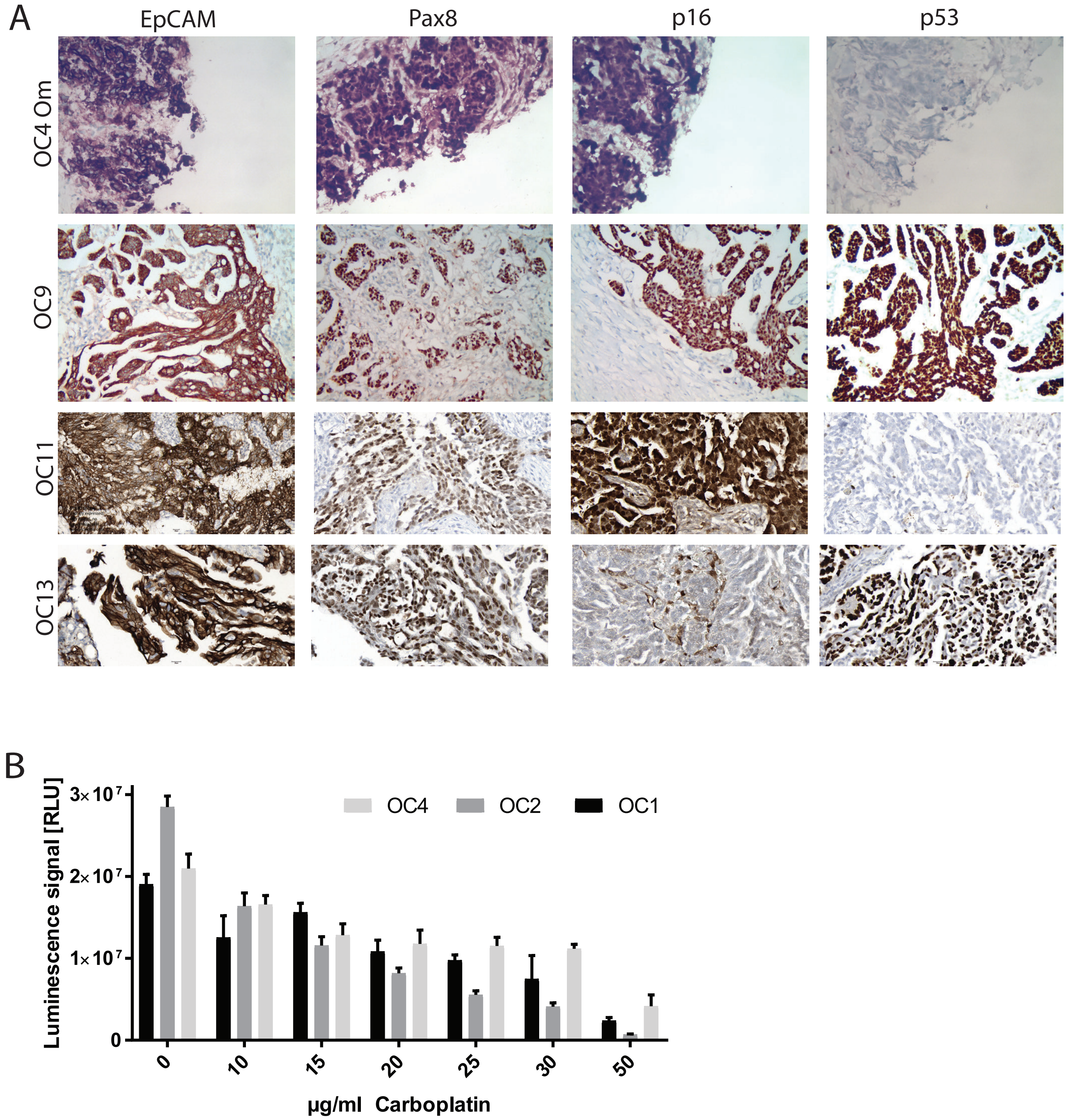

Supplementary Figure 2. Expression profile of HGSOC organoids.

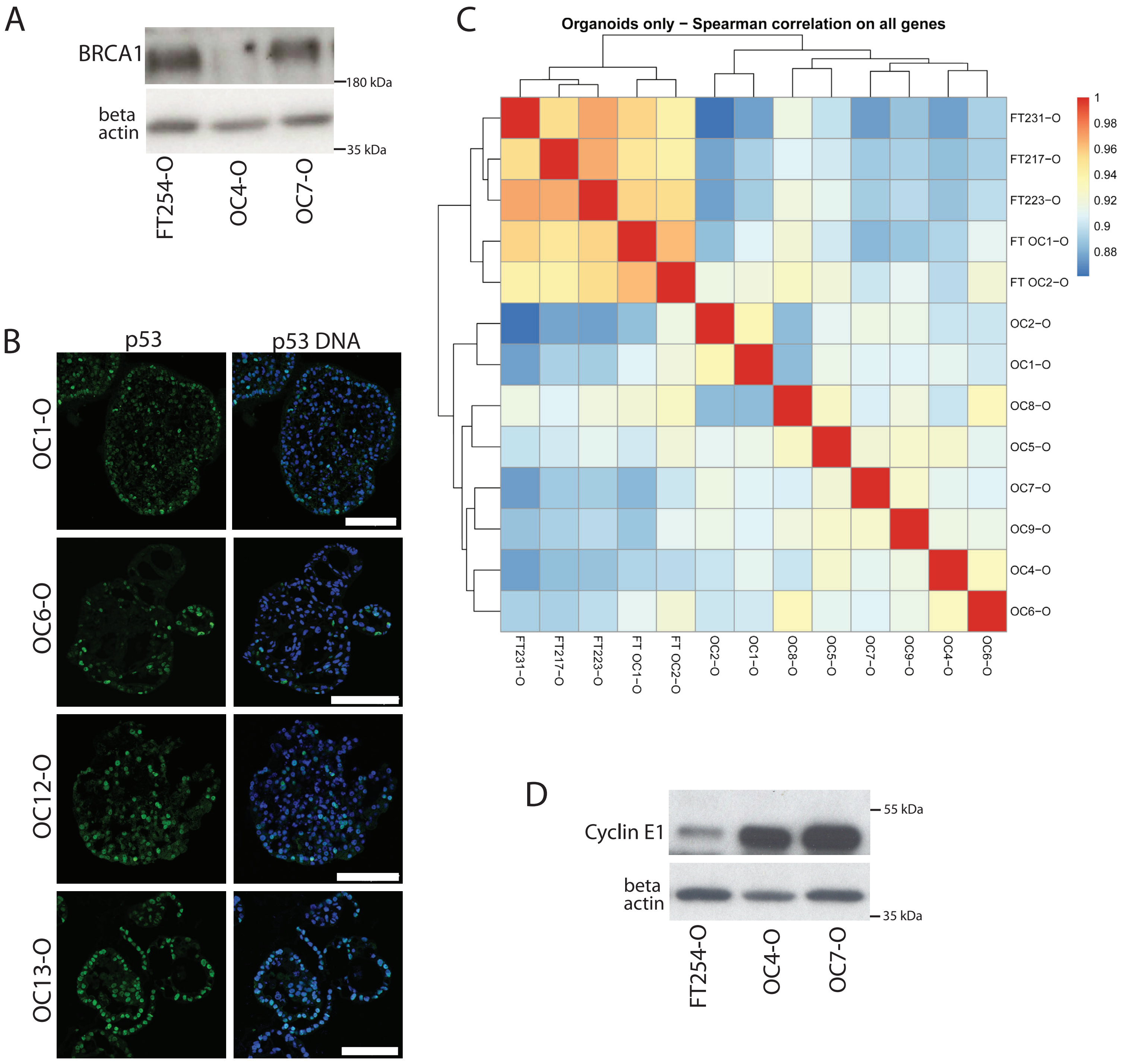

### Supplementary Figure 3. Effects of p53, PTEN and Rb knockdown in FT organoids.

A

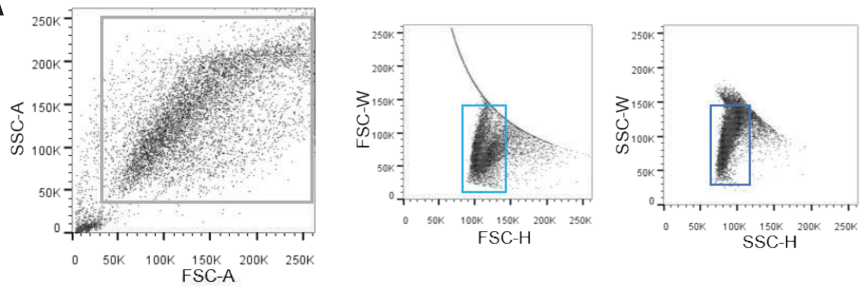

| Population | #Events | %Parent | %Total |
|---|---|---|---|
| All Events | 22,608 | #### | 100.0 |
| P1 | 20,457 | 90.5 | 90.5 |
| P2 | 13,302 | 65.0 | 58.8 |
| P3 | 12,647 | 95.1 | 55.9 |
| P4 | 6,561 | 51.9 | 29.0 |

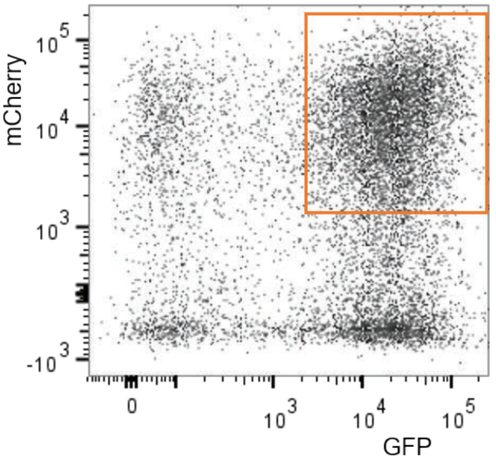

B

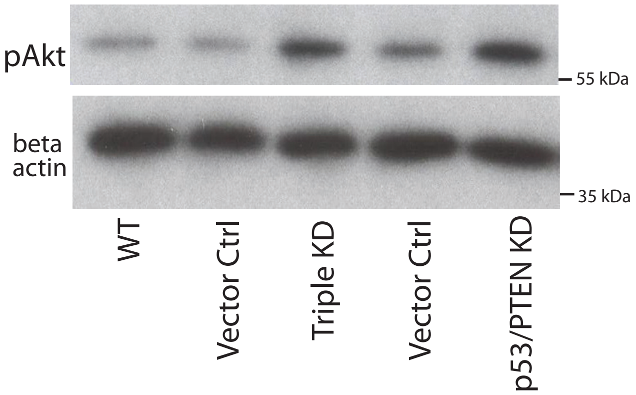

C

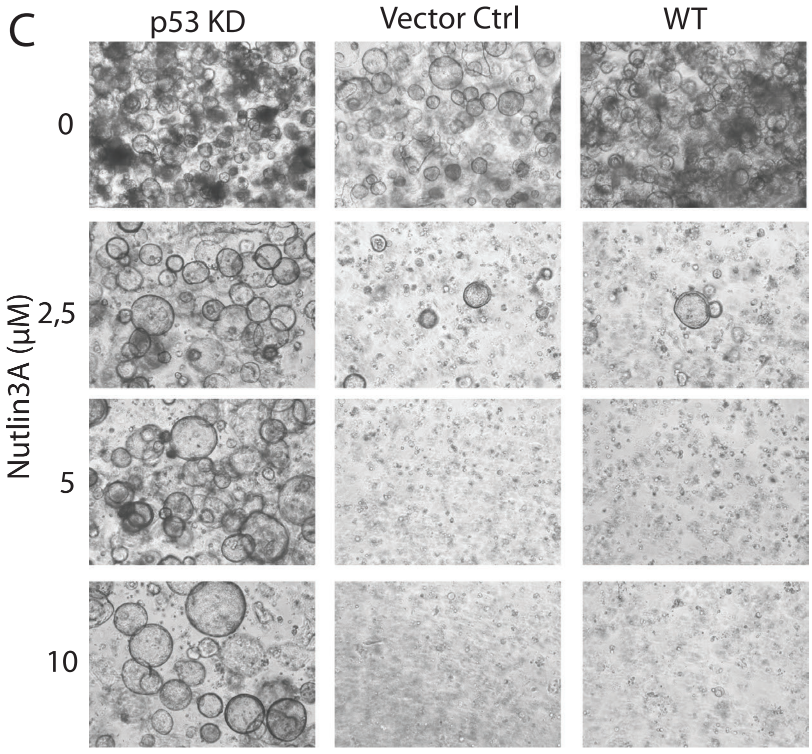

D

| KD organoid line | Passage / time in culture |
|---|---|
| FT219 | p9 / 7 months |
| FT231 | p4 / 3.5 months |
| FT234 | p7 / 5 months |
| FT253 | p5 / 4 months |
| FT254 | p3 / 3 months |
| FT268 | p4 /3 months |
| FT269 | p2 / 1.5 months |

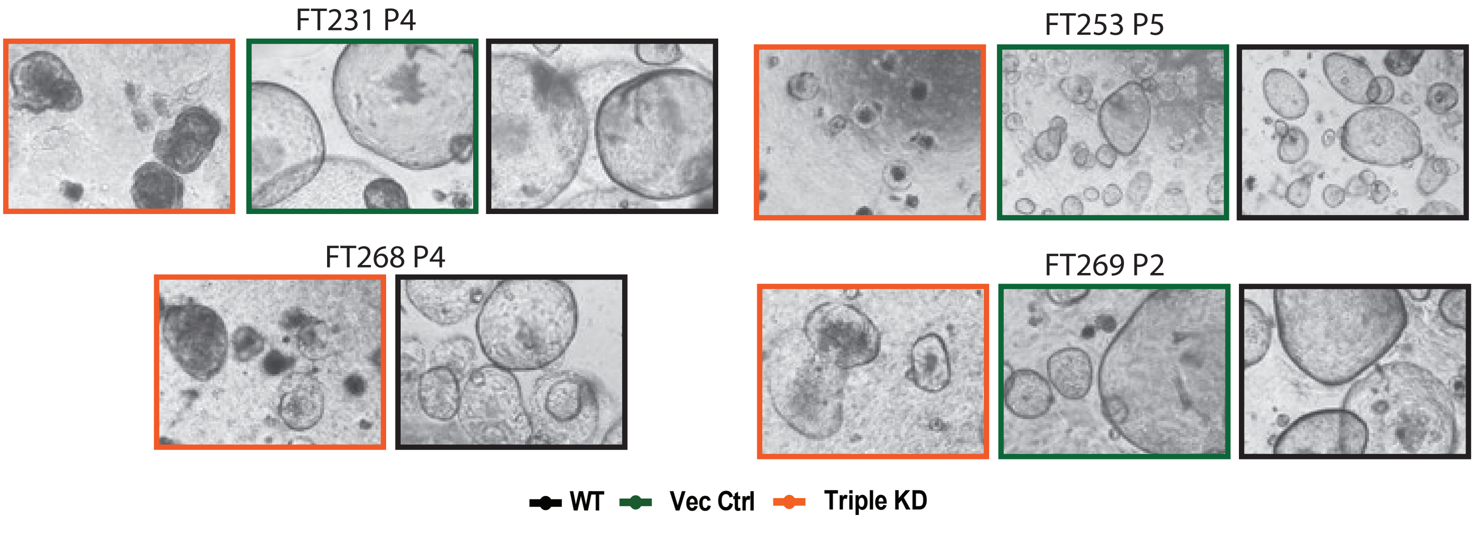

Supplementary Figure 4. Cofirmation of growth arrest, DNA damage and apoptosis as well as activation of Wnt target genes.

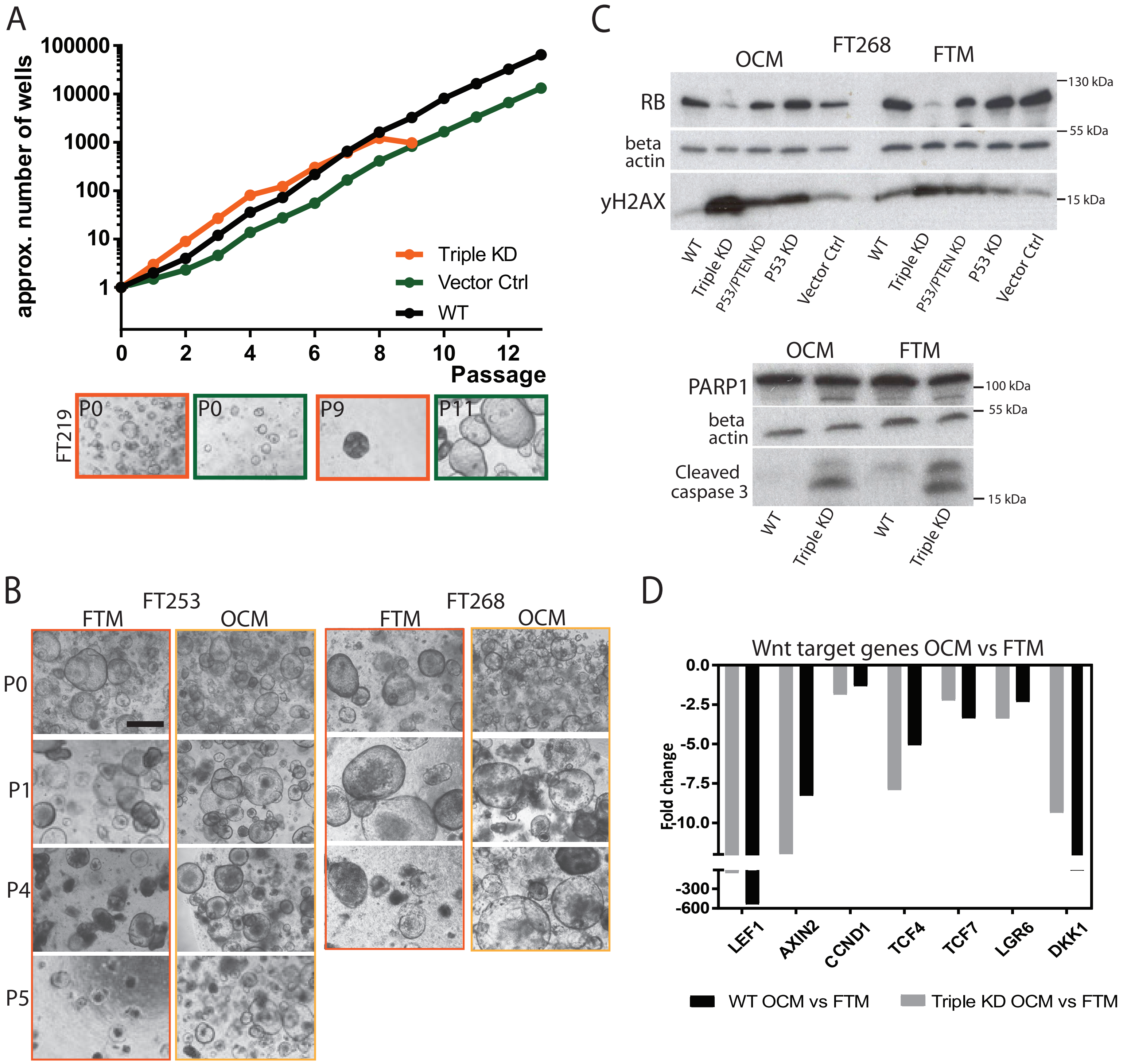

Supplementary Figure 5. HGSOC organoids respond to exogenous Wnt signals with changes in gene expression and growth pattern.

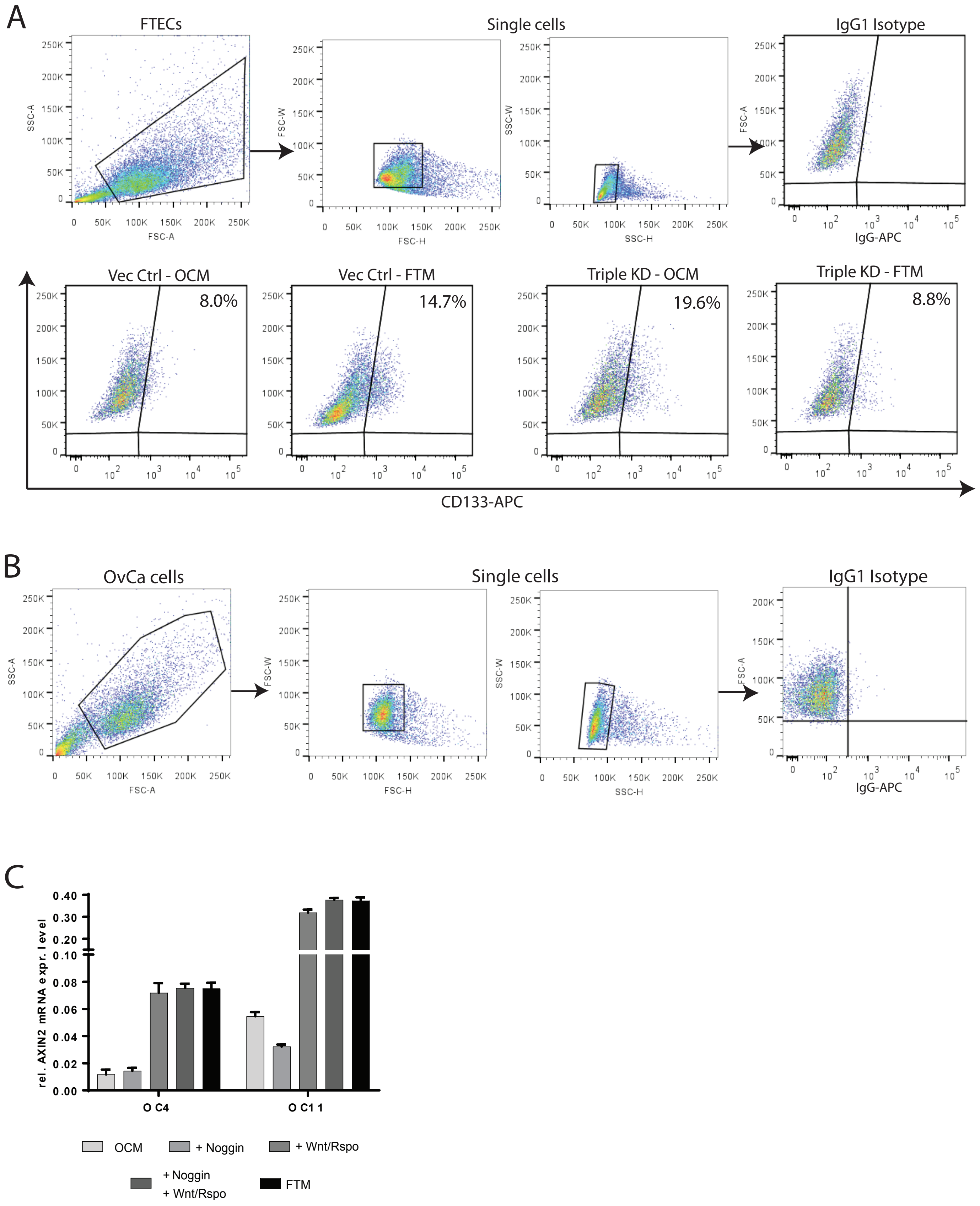

#### Supplementary Table 1 – Patient data of HGSOC samples

| Sample ID | Tumor site | Diagnosis | Histopathology | Age at first disease | FIGO stage | TNM stage | Ascites | Residual mass after surgery | CA125 serum levels | Adjuvant chemotherapy | Cycles | Response |
|---|---|---|---|---|---|---|---|---|---|---|---|---|
| OC1 | Omentum | Primary Ovarian cancer | HGSOC | 74 | IIIC | T3cN1M0 | >500ml | >=2cm | 54 | Carboplatin/Gemcitabine | 10 | CR (Platinum resistant) |
| OC2 | Peritoneum | Primary Ovarian cancer | HGSOC | 69 | IIIC | T3cN1M0 | >500ml | <0.5cm | 900 | - | | |
| OC3 | Peritoneum | Primary Ovarian cancer, Interval surgery | HGSOC | 71 | IVb | T3bN1M1 | No ascites | <0.5cm | 34 | Carboplatin/Paclitaxel/ Avastin | 9 | PR (Platinum resistant) |
| OC4 | Omentum/Peritoneum | Primary Ovarian cancer | HGSOC | 55 | IIIC | T3cN1M0 | >500ml | <1cm | 2800 | Carboplatin/Paclitaxel/ Avastin | 6 | SD (Platinum resistent) |
| OC5 | Omentum/Peritoneum | Primary Peritoneal carcinom | HGSOC | 59 | IIIC | T3cN1M0 | >500ml | No residual mass | 447 | - | | |
| OC6 | Peritoneum | Primary Ovarian Cancer | HGSOC | 29 | IIIb | T3bN1M0 | | <0.5cm | 631 | - | 6 | CR (Platinum sensitiv) |
| OC7 | Peritoneum | Primary Ovarian cancer | HGSOC | 86 | IIIC | T3cN1M0 | | <2cm | | - | | |
| OC8 | Peritoneum | Primary Ovarian cancer | HGSOC | 53 | IIIC | T2bN1M0 | <500ml | No residual mass | 982 | Carboplatin/Paclitaxel/ Avastin or Olaparib | 6 | CR (Platinum sensitiv) |
| OC9 | Peritoneum | Primary Ovarian cancer | HGSOC | 58 | IIIC | T3cN1M0 | >500ml | No residual mass | | Carboplatin/Paclitaxel/ Avastin | 6 | CR (Platinum sensitiv) |
| OC10 | Laparoscopy | Primary Ovarian cancer | HGSOC | 71 | IVb | T3cN0M1 | No ascites | No residual mass | 84 | Carboplatin/Paclitaxel | 6 | CR (Platinum sensitiv) |
| OC11 | Peritoneum | Primary Ovarian cancer | HGSOC | 65 | IVb | T3bN0M1 | >500ml | <2cm | 897 | - | | |
| OC12 | Omentum | Primary Ovarian cancer | HGSOC | 80 | IIIb | T3bN0M0 | <500ml | <0.5cm | 91 | Carboplatin/Paclitaxel/ Avastin | 6 | CR (Platinum sensitiv) |
| OC13 | Omentum | Primary Ovarian cancer | HGSOC | 63 | IVa | T3cN1bM1 | >500ml | 0cm | 1995 | Carboplatin/Paclitaxel/ Avastin | 6 | PR (Platinum resistant) |

| | |
|---|---|
| T- Size of tumor | CR- Complete Response |
| N - Number of lymphnodes | PR- Partial Response |
| M- Number of metastase | SD- Stable Disease |

| FIGO Stage | TNM stage | Description |
|---|---|---|
| I | T1-N0-M0 | Tumor confined to ovaries or fallopian tube(s) |
| IA | T1a-N0-M0 | Tumor limited to 1 ovary (capsule intact) or fallopian tube; no tumor on ovarian or fallopian tube surface; no malignant cells in the ascites or peritoneal washings |
| IB | T1b-N0-M0 | Tumor limited to both ovaries (capsules intact) or fallopian tubes; no tumor on ovarian or fallopian tube surface; no malignant cells in the ascites or peritoneal washings |
| IC1 | T1c1-N0-M0 | Tumor limited to 1 or both ovaries or fallopian tubes, with surgical spill |
| IC2 | T1c2-N0-M0 | Tumor limited to 1 or both ovaries or fallopian tubes, with capsule ruptured before surgery or tumor on ovarian or fallopian tube surface |
| IC3 | T1c3-N0-M0 | Tumor limited to 1 or both ovaries or fallopian tubes, with malignant cells in the ascites or peritoneal washings |
| II | T2-N0-M0 | Tumor involves 1 or both ovaries or fallopian tubes with pelvic extension (below pelvic brim) or peritoneal cancer |
| IIA | T2a-N0-M0 | Extension and/or implants on uterus and/or fallopian tubes and/or ovaries |
| IIB | T2b-N0-M0 | Extension to other pelvic intraperitoneal tissues |
| III | T1/T2-N1-M0 | Tumor involves 1 or both ovaries or fallopian tubes, or peritoneal cancer, with cytologically or histologically confirmed spread to the peritoneum outside the pelvis and/or metastasis to the retroperitoneal lym |
| IIIA1 | T3a2-N0/N1-M0 | Positive retroperitoneal lymph nodes only (cytologically or histologically proven) |
| IIIA2 | T3a2-N0/N1-M0 | Microscopic extrapelvic (above the pelvic brim) peritoneal involvement with or without positive retroperitoneal lymph nodes |
| IIIB | T3b-N0/N1-M0 | Macroscopic peritoneal metastasis beyond the pelvis up to 2 cm in greatest dimension, with or without metastasis to the retroperitoneal lymph nodes |
| IIIC | T3c-N0/N1-M0 | Macroscopic peritoneal metastasis beyond the pelvis more than 2 cm in greatest dimension, with or without metastasis to the retroperitoneal lymph nodes |
| IV | Any T, any N, M1 | Distant metastasis excluding peritoneal metastases |
| IVA | Any T, any N, M1 | Pleural effusion with positive cytology |
| IVB | Any T, any N, M1 | Parenchymal metastases and metastases to extra-abdominal organs |

**According to the FIGO Cancer Report 2018**

#### Supplementary Table 2 – HGSOC Medium composition

| Sample | Deposit | Medium | Time in culture |
|---|---|---|---|
| OC1 | Om | Basic +EFR | p17 (12.5 months) |
| OC2 | Peri | Basic +EFR | p20 (13 months) |
| OC3 | Peri | Basic +E | p15 |
| OC4-1 | Peri | Basic +E | p26 (16 months) |
| OC4-2 | Om | Basic +EFNR | p26 |
| OC5-1 | Peri | Basic +E | p8 |
| OC5-2 | Om | Basic +E | p8 |
| OC6 | Peri | Basic +E | p8 |
| OC7 | Peri | Basic +EB | p26 (17 months) |
| OC8 | Peri | Basic +EFN | p10; limited expansion in later passage |
| OC9 | Peri | Basic +E | p6 |
| OC10 | L | Basic +EB | p23 (14 months) |
| OC11 | Peri | Basic +EB | p23 (12.5 months) |
| OC12 | Peri | Basic +EB | p17 |
| OC13 | Peri | Basic +EB | p17 |

| | |
|---|---|
| (blue) | active BMP signaling, low Wnt |
| (green) | active BMP, high Wnt |
| (yellow) | BMP inhibition |

| | | |
|---|---|---|
| Om - Omentum | | p - Passage |
| Peri - Peritoneum | | |
| L - Laparoscopy | | |
| Basic = Nicotinamide, B27, N2, ROCKi, TGF-β inhibitor | | |
| Fallopian tube medium = Basic +EGF +Noggin +FGF +RSPO1 +Wnt3a | | |
| Ovarian Cancer Medium = Basic +EGF +BMP2 | | |
| B | BMP2 | |
| E | EGF | |
| F | FGF | |
| N | Noggin | |
| R | R-Spondin1 | |

#### Supplementary Table 3 – Target genes for sequencing

| Target genes | |
|---|---|
| ABL1 | KIT |
| ALK | KRAS |
| APC | LEF1 |
| ARID1A | LRP5 |
| ATM | MAML1 |
| ATR | MAML2 |
| BACH1 | MAML3 |
| BARD1 | MAP3K7 |
| BRAF | MAPK2K1 |
| BRCA1 | MET |
| BRCA2 | MLH1 |
| BRD4 | MMP7 |
| BRF2 | MPL |
| BRIP1 | MSH2 |
| CCNE1 | MSH6 |
| CDH1 | MYC |
| CDK12 | NBN |
| CDKN2A | NF1 |
| CHD8 | NFAT5 |
| CHEK2 | NFATC3 |
| CREBBP | NOTCH1 |
| CSF1R | NOTCH3 |
| CSMD3 | NPM1 |
| CSNK2A1 | NRAS |
| CTBP2 | PALB2 |
| CTNNB1 | PDGFRA |
| CUL1 | PDGFRB |
| DAAM2 | PIK3CA |
| DDR2 | PIK3R1 |
| DSN1 | PMS2 |
| EGFR | PPP2R1A |
| EMSY | PPP3P1 |
| EP300 | PRICKLE1 |
| ERBB2 | PRICKLE2 |
| ERBB4 | PRKACG |
| EZH2 | PTEN |
| FANCA | PTPN11 |
| FANCI | RAD50 |
| FAT3 | RAD51B |
| FBXW7 | RAD51C |
| FGFR1 | RAD51D |
| FGFR2 | RAD51L1 |
| FGFR3 | RAD51L3 |
| FLT3 | RAD52 |
| FZD1 | RAD54L |
| FZD3 | RB1 |
| GABRA6 | RET |
| GNA11 | ROCK1 |
| GNAQ | ROCK2 |
| GNAS | SENP2 |
| HNF1A | SMAD4 |
| HRAS | SMARCB1 |
| IDH1 | SMO |
| IDH2 | SRC |
| IGF1R | STK11 |
| JAK1 | TCF7L2 |
| JAK2 | TGFBR2 |
| JAK3 | TP53 |
| KDR | TPX2 |
| | VANGL1 |
| | VANGL2 |
| | VHL |

### Supplementary Table 4 – Allelic frequencies of mutations identified by targeted sequencing in tissue and organoids

| Gene | Mutation (HGVS.c) | Effect | rare in healthy | OC2 - T | OC2 - O | OC4 P - T | OC4 P - O | OC4 Om - T | OC4 Om - O | OC5 - T | OC5 - O | OC6 - T | OC6 - O | OC7 - T | OC7 - O | OC8 - T | OC8 - O | OC10 - T | OC10 - O | OC11 - T | OC11 - O |
|---|---|---|---|---|---|---|---|---|---|---|---|---|---|---|---|---|---|---|---|---|---|
| **ATM** | c.4534G>A | missense_variant | yes, <1% of population | | | | | | | | | | | | | | | | | 0,452 | 0,514 |
| **ATM** | c.2119T>C | missense_variant | yes, <1% of population | 0,461 | 0,491 | | | | | | | | | | | | | | | | |
| **ATR** | c.1517A>G | missense_variant | yes, <1% of population | | | | | | | | | 0,603 | 0,627 | | | | | | | | |
| **ATR** | c.4576A>G | missense_variant | yes, <1% of population | | | | | | | | | 0,589 | 0,636 | | | | | | | | |
| **BRCA1** | c.4900A>G | missense_variant | no | | | | | | | 0,774 | 0,987 | | | 0,988 | 0,993 | 0,24 | | 0,911 | 0,984 | | |
| **BRCA1** | c.3113A>G | missense_variant | no | | | | | | | 0,784 | 0,991 | | | 0,982 | 0,989 | 0,208 | | 0,909 | 0,992 | | |
| **BRCA1** | c.2612C>T | missense_variant | no | | | | | | | 0,774 | 0,998 | | | 0,997 | 0,998 | 0,207 | | 0,921 | 0,999 | | |
| **CREBBP** | c.5933A>G | missense_variant | yes, <1% of population | 0,377 | 0,459 | | | | | | | | | | | | | | | | |
| **EGFR** | c.1562G>A | missense_variant | no | | | | | | | 0,992 | 0,988 | | | | | 0,36 | 0,303 | | | | |
| **EP300** | c.6288T>G | missense_variant | yes, <1% of population | 0,547 | 0,486 | | | | | | | | | | | | | | | | |
| **ERBB2** | c.1963A>G | missense_variant | no | 0,602 | 0,473 | | | | | | | | | | | 0,763 | 0,995 | | | | |
| **FANCA** | c.1238G>T | missense_variant | yes, <1% of population | | | | | | | | | | | 0,453 | 0,465 | | | | | | |
| **FAT3** | c.10552G>T | missense_variant | no | 0,515 | 0,479 | | | | | 0,997 | 0,997 | | | | | 0,416 | 0,47 | 0,548 | 0,353 | | |
| **FZD9** | c.434G>A | missense_variant | yes, <1% of population | | | | | | | | | | | | | | | | | 0,355 | 0,28 |
| **KIT** | c.1621A>C | missense_variant | no | | | | | | | 0,497 | 0,535 | 0,481 | 0,511 | | | | | | | | |
| **LRP5** | c.2335G>A | missense_variant | yes, <1% of population | | | | | | | | | | | | | 0,485 | 0,495 | | | | |
| **MAML1** | c.1526T>G | missense_variant | yes, <1% of population | | | | | | | | | 0,485 | 0,479 | | | | | | | | |
| **MLH1** | c.655A>G | missense_variant | no | | | 0,512 | 0,498 | 0,516 | 0,75 | 0,62 | 0,763 | 0,602 | 0,631 | 0,568 | 0,341 | | | 0,48 | 0,487 | | |
| **MSH6** | c.472C>T | missense_variant | no | | | | | | | | | 0,861 | 0,966 | 0,574 | 0,626 | | | 0,454 | 0,694 | | |
| **NOTCH3** | c.3399C>A | missense_variant | no | 0,502 | 0,482 | | | | | | | | | | | | | | | | |
| **PTEN** | c.464A>G | missense_variant | yes, <1% of population | | | | | | | | | | | | | | 0,454 | | | | |
| **ROCK2** | c.1313A>G | missense_variant | yes, <1% of population | | | | | | | | | | | 0,579 | 0,673 | | | | | | |
| **ROCK2** | c.1292C>A | missense_variant | no | 0,995 | 0,999 | 0,491 | 0,481 | 0,501 | 0,242 | 0,997 | 0,997 | | | 0,409 | 0,322 | 0,998 | 0,997 | 0,989 | 0,996 | 0,29 | 0,318 |
| **TCF7L2** | c.1535C>G | missense_variant | yes, <1% of population | 0,624 | 0,591 | | | | | | | | | | | | | | | | |
| **TP53** | c.524G>A | missense_variant | yes, <1% of population | | | | | | | | | | | | | | | | 0,986 | | |
| **TP53** | c.501dupG | frameshift_variant | yes, <1% of population | | | | | | | 0,541 | 0,958 | | | | | | | | | | |
| **TP53** | c.283delT | frameshift_variant | yes, <1% of population | | | | | | | | | | | 0,304 | 0,956 | | | | | | |
| **TP53** | c.660T>G | stop_gained | yes, <1% of population | | | | | | | | | | | | | | | | | 0,58 | 0,994 |
| **TP53** | c.993+1G>C | splice_donor_variant& | yes, <1% of population | 0,22 | | | | | | | | | | | | | | | | | |
| **TP53** | c.517G>A | missense_variant | yes, <1% of population | | | | | | | | | 0,711 | 0,983 | | | | | | | | |
| **TP53** | c.560-1G>A | splice_acceptor_varia | yes, <1% of population | | | 0,418 | 0,993 | 0,693 | 0,995 | | | | | | | | | | | | |
| **TP53** | c.817C>T | missense_variant | yes, <1% of population | | | | | | | | | | | | | 0,515 | 0,987 | | | | |

| |
|---|
| O - Organoid |
| T - Tissue |

#### Supplementary Table 5 – Protein expression in OC organoids

| | p53 | CyclinE1 | BRCA1 | RB |
|---|---|---|---|---|
| OC1-O | gain | not tested | not tested | not tested |
| OC2-O | complete loss | gain | partial loss/down-reg. | unchanged |
| OC3-O | gain | unchanged | partial loss/down-reg. | unchanged |
| OC4-O | complete loss | gain | partial loss/down-reg. | unchanged |
| OC5-O | complete loss | not tested | not tested | not tested |
| OC6-O | gain | not tested | not tested | not tested |
| OC7-O | complete loss | gain | unchanged | unchanged |
| OC8-O | gain | not tested | not tested | not tested |
| OC9-O | gain | not tested | not tested | not tested |
| OC10-O | gain | gain | unchanged | unchanged |
| OC11-O | complete loss | gain | unchanged | unchanged |
| OC12-O | gain | unchanged | unchanged | unchanged |
| OC13-O | gain | unchanged | unchanged | unchanged |

| | |
|---|---|
| | unchanged |
| | gain |
| | partial loss/down-reg. |
| | complete loss |
| | not tested |

#### Supplementary Table 6 – Microarray data of Wnt target genes

| | WT FTM vs Triple KD FTM | | | | | Triple KD FTM vs Triple KD OCM | | | |
|---|---|---|---|---|---|---|---|---|---|
| **Gene** | **Fold Change** | **P-value** | **Intensity1** | **Intensity2** | | **Fold Change** | **P-value** | **Intensity1** | **Intensity2** |
| LEF1 | -6,22 | 1,68E-27 | 4824 | 710 | | -54,44 | 7,93E-20 | 624 | 9 |
| AXIN2 | -1,41 | 8,62E-07 | 142 | 101 | | -6,15 | 0 | 97 | 16 |
| CCND1 | -1,76 | 0,00189 | 4362 | 2443 | | -1,86 | 0,00001 | 2685 | 1477 |
| TCF4 | -1,41 | 0,00501 | 3034 | 2087 | | -7,90 | 0 | 1649 | 210 |
| DKK3 | 2,94 | 0,00 | 14,52 | 39,20 | | 2,86 | 0,00 | 36,79 | 104,88 |
| KREMEN2 | 3,44 | 1,24E-31 | 43 | 145 | | -1,26 | 0,00298 | 131 | 103 |
